# Cellular toxicity of iHAP1 and DT-061 does not occur through PP2A-B56 targeting

**DOI:** 10.1101/2021.07.08.451586

**Authors:** Gianmatteo Vit, Joana Duro, Girish Rajendraprasad, Emil P.T. Hertz, Lya Katrine Kauffeldt Holland, Melanie Bianca Weisser, Brennan C. McEwan, Blanca Lopez-Mendez, Guillermo Montoya, Niels Mailand, Kenji Maeda, Arminja Kettenbach, Marin Barisic, Jakob Nilsson

**Author notes:** Equal contribution (Gianmatteo Vit, Joana Duro, Girish Rajendraprasad).

## Abstract

PP2A is an abundant phosphoprotein phosphatase that acts as a tumor suppressor. For this reason, compounds able to activate PP2A are attractive anticancer agents. The small molecule compounds iHAP1 and DT-061 have recently been reported by Leonard et al. (2020) and Morita et al. (2020) in *Cell* to selectively stabilize specific PP2A-B56 complexes which mediate cell killing. Here, we show that this is not the case and question key findings in these papers. Through genome wide CRISPR-Cas9 screens, we uncover the biological pathways targeted by these compounds. We find that iHAP1 directly blocks microtubule assembly both *in vitro* and *in vivo* and thus acts as a microtubule poison. In contrast, DT-061 disrupts both the Golgi apparatus and the endoplasmic reticulum and we directly visualize DT-061 in cytoplasmic granules that co-localize with Golgi markers. Our work demonstrates that iHAP1 and DT-061 cannot be used for dissecting PP2A-B56 biology.

## Introduction

PP2A is an abundant phosphoprotein phosphatase (PPP) and accounts for much of the serine/threonine phosphatase activity in eukaryotic cells, hereby suppressing multiple oncogenic kinase pathways [1–4]. The PP2A holoenzyme consists of a catalytic subunit (PP2AC, A and B isoforms), a scaffold subunit (PPP2R1, A and B isoforms) and one of four regulatory B-subunits (B55α-δ, B56α-ε, PR72/PR130 or STRN1-4, multiple isoforms of each B subunit) [5–7]. The B subunits confer specificity to PP2A holoenzymes by directly binding to substrates or substrate specifiers [6, 8]. Given that PP2A activity is suppressed in cancers either through overexpression of inhibitory proteins or through mutations in PP2A components, there is an interest in developing compounds that can reactivate PP2A for cancer treatment [4, 9].

Several compounds based on a phenothiazine or phenoxazine scaffold, which are both tricyclic heterocycle structures and include iHAP1 and DT-061, have been shown to modulate PP2A holoenzyme stoichiometry, thus mediating their anticancer activity [10–12]. In a screen for FDA-approved compounds that could kill T-cell acute lymphoblastic leukemia (T-ALL) cells, the antipsychotic drug perphenazine could be identified [12]. Through mass spectrometry-based approaches the target of perphenazine was found to be PPP2R1A. In a parallel work, tricyclic heterocyclic compounds that affected FoxO1 localization were identified and reengineered to remove their effects on the central nervous system, making them more cancer specific compounds [13]. This reengineering let to the identification of several compounds (small molecule activators of PP2A, SMAPs), including DT-061, that was found to bind PPP2R1A with a Kd of 235 nM, and the binding site mapped to amino acids 194-198 of PPP2R1A. In *in vitro* assays, 20 μM DT-061 was found to activate PP2A-B56γ and the PPP2R1A-PP2AC complex by 20-30% [14].

In two recent back-to-back publications in *Cell*, perphenazine and DT-061 discoveries were extended [10, 11]. Morita et al. (2020) improved the perphenazine scaffold to remove, among others, off target effects on microtubules to generate a novel compound named iHAP1 (Fig. 1A) [11]. The identification of iHAP1 was in part based on the screening of a compound library by adding compounds to cells for 3 hours, followed by purification of affinity tagged PP2AC coupled with activity testing. It was found that iHAP1 specifically stabilizes PP2A-B56ε in leukemic and Kelly cells and that this causes a strong arrest in prometaphase through the selective dephosphorylation of MYBL2 by PP2A-B56ε. DT-061 mechanism of action was also investigated by Morita et al. (2020) using a similar approach, and they found that the target was (is?) PP2A-B55α [11]. These results contrast with Leonard et al. (2020) in the same issue of *Cell* that carried out a structure-function analysis of DT-061 and identified PP2A-B56α as the target of DT-061 [10]. A PP2A-B56α-DT-061 complex was analyzed by cryo-EM and the analysis revealed that DT-061 interacts with an interface formed by all the three subunits of the PP2A-B56α holoenzyme. This interface is close to the very C-terminal flexible tail of the PP2AC subunit, and specific residues in B56α contact DT-061, explaining the selective binding to PP2A-B56α and not the other B56 isoforms.

**Figure 1.**
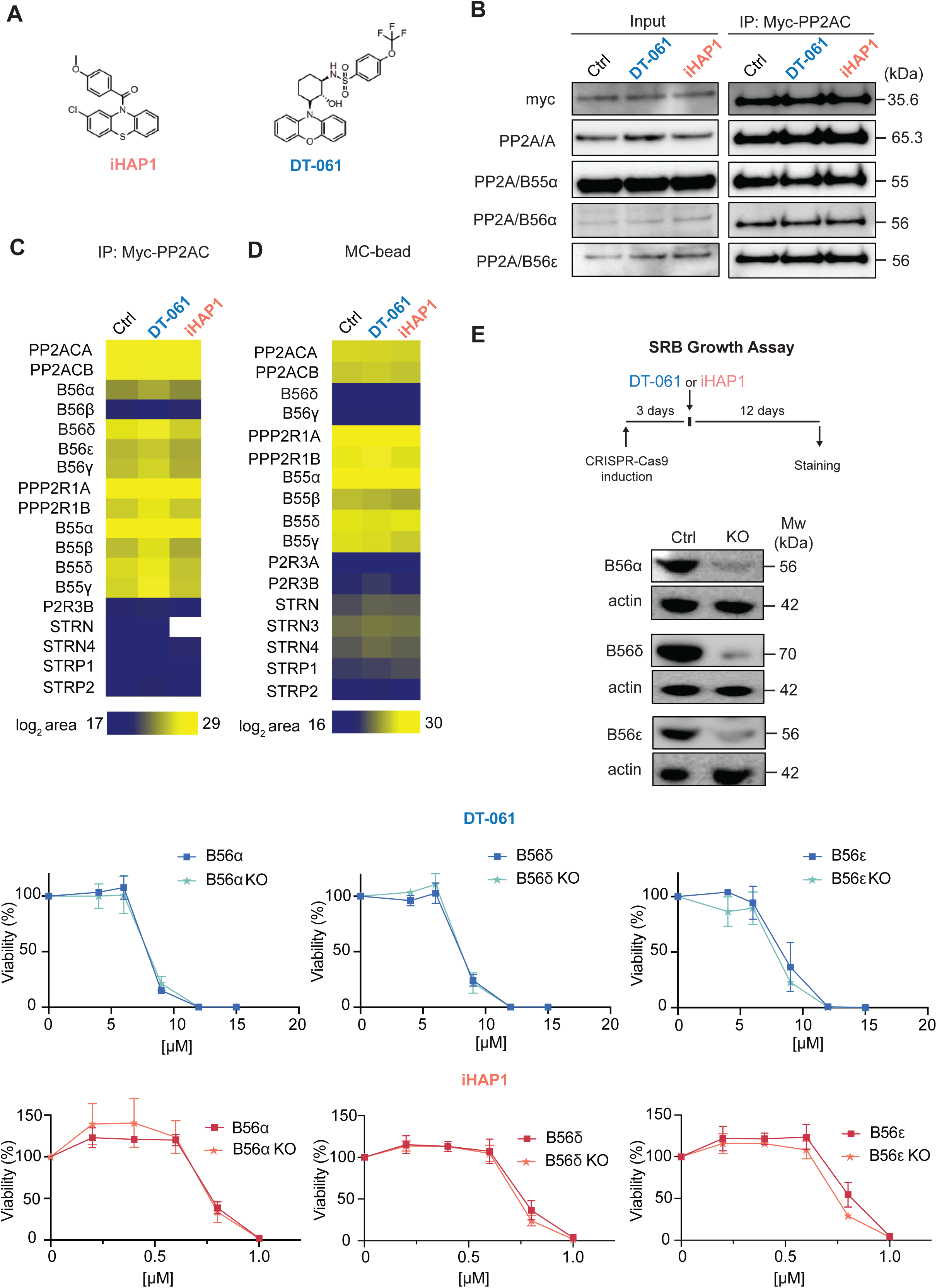
iHAP1 and DT-061 toxicity is not affected by knockdown of specific B56 subunits. A) Chemical structure of iHAP1 and DT-061. B) A stable HEK293 cell line expressing myc- PP2AC was treated with DMSO, DT-061 or iHAP1 for 30 minutes and myc-PP2AC affinity purified. The binding of the indicated proteins was analysed by Western blotting and C) by quantitative mass spectrometry. D) HEK-293T cells were incubated with DMSO, DT-061 or iHAP1 for 30 minutes and protein lysates were incubated with a microcystin affinity column to capture PPP complexes. Bound complexes were analysed by mass spectrometry to identify PP2A components. E) Doxycyclin-inducible CRISPR-Cas9 HeLa cell lines allows for depletion of specific B56 subunits. Western blotting shows the removal of the indicated B56 subunits 3 days after addition of doxycycline. Actin was used as loading control. Growth assays in the presence of the indicated concentrations of DT-061 or iHAP1 measured after 12 days of addition of compound. Mean and SD of three independent experiments is shown.

Based on our inability to detect any effect of iHAP1 and DT-061 on PP2A holoenzyme activity in *in vitro* reconstituted systems and a failure to reproduce specific binding of these compounds to PPP2R1A by ITC and NMR (Fig. S1-S4), we decided to investigate these compounds carefully.

## Results

### The cellular toxicity of iHAP1 and DT-061 is not dependent on specific B56 subunits

Given that we could not confirm an effect of the compounds on PP2A activity *in vitro*, we set out to investigate the impact on PP2A holoenzyme formation in cells. Incubation with iHAP1 or DT-061 has been shown to affect PP2A holoenzyme stoichiometries in cells through selective stabilization of PP2A-Β56ε and PP2A-B56α, respectively [10, 11]. We therefore treated HEK-293T cells stably expressing myc-tagged PP2AC with 2 μM iHAP1 or 20 μM DT-061 for 30 minutes, a timeframe sufficient to induce clear cellular effects (see below), and immunopurified myc-PP2AC. We did not detect any effect on the binding of tested B subunits with PP2AC by Western blotting or B56 subunits detected by quantitative mass spectrometry analysis (Fig. 1B-C, Table S1). In a complementary approach, we incubated lysates from untreated or iHAP1/DT-061 treated HEK-293T cells with microcystin coupled to beads and analysed by mass spectrometry the composition of endogenous PPP components which were pulled down by microcystin [15] (Fig. 1D, Table S2). If a certain PP2A holoenzyme is stabilized by the compound, one would detect this in the unbiased MS interactome analysis. Again, we observed no statistically significant changes in the detection of B56 or B55 subunits (p-value < 0.05, log2 ratio > 1).

To explore if the cellular toxicity of DT-061 and iHAP1 is dependent on specific PP2A-B56 complexes, an inducible CRISPR-Cas9 system was used[16]. We induced the expression of gRNAs to B56α, B56δ and B56ε in HeLa cells stably expressing Cas9, which resulted in penetrant depletion of these B56 subunits (Fig. 1E). We then monitored the effect of iHAP1 and DT-061 on cell viability and determined LC50. We observed no effect on LC50 values when the proposed targets of iHAP1 and DT-061 were removed by CRISPR, arguing that the toxicity of these compounds is unrelated to the stabilization and activation of specific PP2A-B56 complexes.

Based on these results, we reinvestigated the cryo-EM structure of the proposed PP2A-B56α-DT-061 complex published in Leonard et al. (2020). This revealed that the assignment of density to DT-061 is not unambiguous at the given intermediate resolution of 3.6 Å (Fig. S5). The density assigned to DT-061 does not fully embed the tricyclic ring of DT-061 and might as well be ascribed to residues from the adjacent C-terminal tail of PP2AC, which is known to be flexible and therefore only partially modelled in this study as well as in several other PP2A structures available [17–19]. Structures of unliganded and liganded complexes are needed to fully establish that the assigned density can be attributed to DT-061.

Collectively, our analysis demonstrates that DT-061 and iHAP1 do not affect specific PP2A-B56 complexes to mediate their cellular toxicity.

### Genome-wide CRISPR-Cas9 screens identify cellular pathways sensitizing cells to iHAP1 and DT-061

Since iHAP1 and DT-061 could still be relevant therapeutic compounds for cancer treatment, we set out to identify their mechanisms of cellular toxicity. Genome-scale CRISPR dropout screens are powerful tools to uncover genetic vulnerabilities to chemical compounds and can provide insight into the molecular pathways affected by these drugs [20–23]. To identify genes whose inactivation causes increased sensitivity to iHAP1 or DT-061, we transduced the RPE1-hTERT P53-/- Flag-Cas9 cell line [24] with the TKOv3 lentiviral library of single-guide (sg) RNAs targeting 18,053 human genes and monitored the depletion of sgRNAs after 12 days of low-dose (LD20) drug treatment compared to control treated cells (Fig. 2A and Fig. S6A) [24]. A normalized depletion score was calculated for each gene using the DrugZ software [25]. This revealed striking differences in the chemogenetic profile of DT-061 and iHAP1 (Fig. 2A, Table S3). Firstly, we identified no genes related to PP2A biology in our top ranked hits for either of the compounds. Depletion of genes related to Golgi and endoplasmic reticulum (ER) function, such as components of the NatC complex and TRAPP complex, resulted in increased sensitivity to DT-061, while known mitotic regulators induced the strongest sensitization to iHAP1 (Fig. 2A).

**Figure 2.**
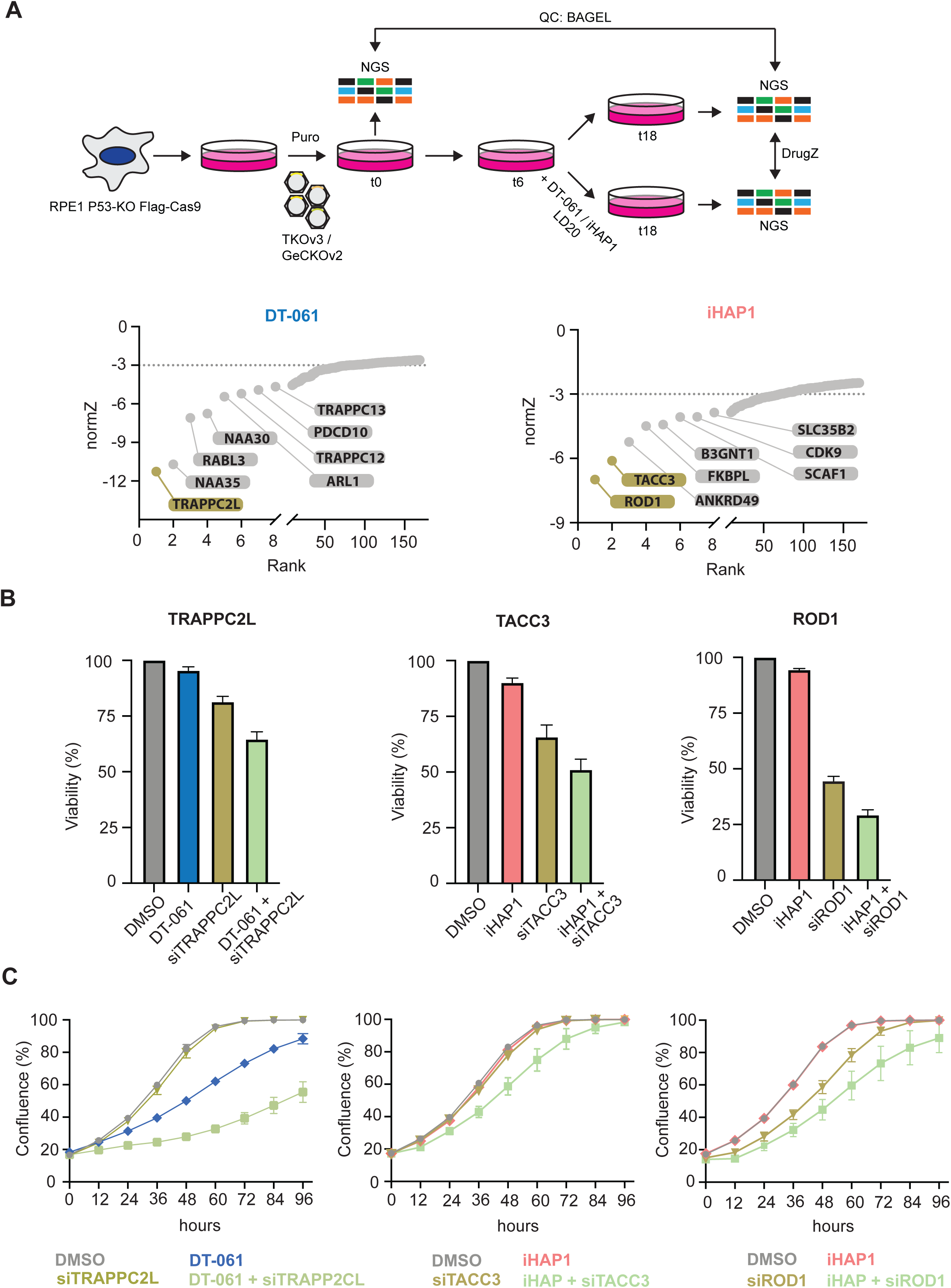
Genome-wide CRISPR screen establishes sensitizers to iHAP1 and DT-061. A) Schematic of the experimental setup and results of CRISPR screen. Genes were ranked based on their normZ score. The top hits which were further analysed are highlighted. B) Validation of CRISPR screen results using RNAi depletion followed by SRB growth assay. Mean and SD of three independent experiments is shown. C) Incucyte growth assay after RNAi depletion of indicated proteins alone or in combination with DT-061 or iHAP1 treatment. B-C) Mean and SD of three independent experiments is shown.

We validated the top hits from our CRISPR screens by an independent method. TRAPPC2L, Rod1, and TACC3 were depleted by RNAi in the RPE1-hTERT P53-/- Flag-Cas9 cell line, and sensitivity to iHAP1 and DT-061 was determined using two different cell growth assays (Fig. 2B-C and Fig. S6B). Consistent with the CRISPR screen, the depletion of TRAPPC2L resulted in increased sensitivity to DT-061, while depletion of Rod1 and TACC3 resulted in sensitization to iHAP1. We further validated the chemogenic interaction between Rod1 and iHAP1 by monitoring cell growth of a previously generated HeLa Rod1-/- cell line [26]. As for the RPE1-hTERT P53-/- background, deletion of Rod1 in HeLa cells caused marked sensitization to iHAP1 (Fig. S6C-D).

Collectively, the difference in chemogenic profile between the two drugs argues that the molecular targets of DT-061 and iHAP1 must be distinct. Based on the chemogenic profiles, we expect iHAP1 to affect mitotic processes while DT-061 affects Golgi/ER function. We decided to explore this in more detail.

### iHAP1 is a microtubule poison

TACC3 is a microtubule stabilizer during mitosis and an important determinant of the response to microtubule poisons [27, 28]. The fact that TACC3 depletion increased the sensitivity to iHAP1 raised the possibility that iHAP1 might affect microtubules directly. Phenothiazine derived compounds can disturb microtubule assembly; a previous study had shown that iHAP1 is a microtubule poison [29] while Morita et al. (2020) reported no effect on microtubule polymerization *in vitro*.

In light of this discrepancy, we tested if iHAP1 has a direct effect on microtubule dynamics. We first conducted *in vitro* microtubule polymerization assays and analysed the effect of iHAP1 and DT-061 on this. While DT-061 had no effect on microtubule polymerization, we observed a clear effect of iHAP1 at 2 μM, the usual concentration in cellular assays. At 5 µM iHAP1, microtubule polymerization was fully blocked, similar to nocodazole treatment, a well-established microtubule poison (Fig. 3A). We were thus unable to confirm a key claim in Morita et al. (2020).

**Figure 3.**
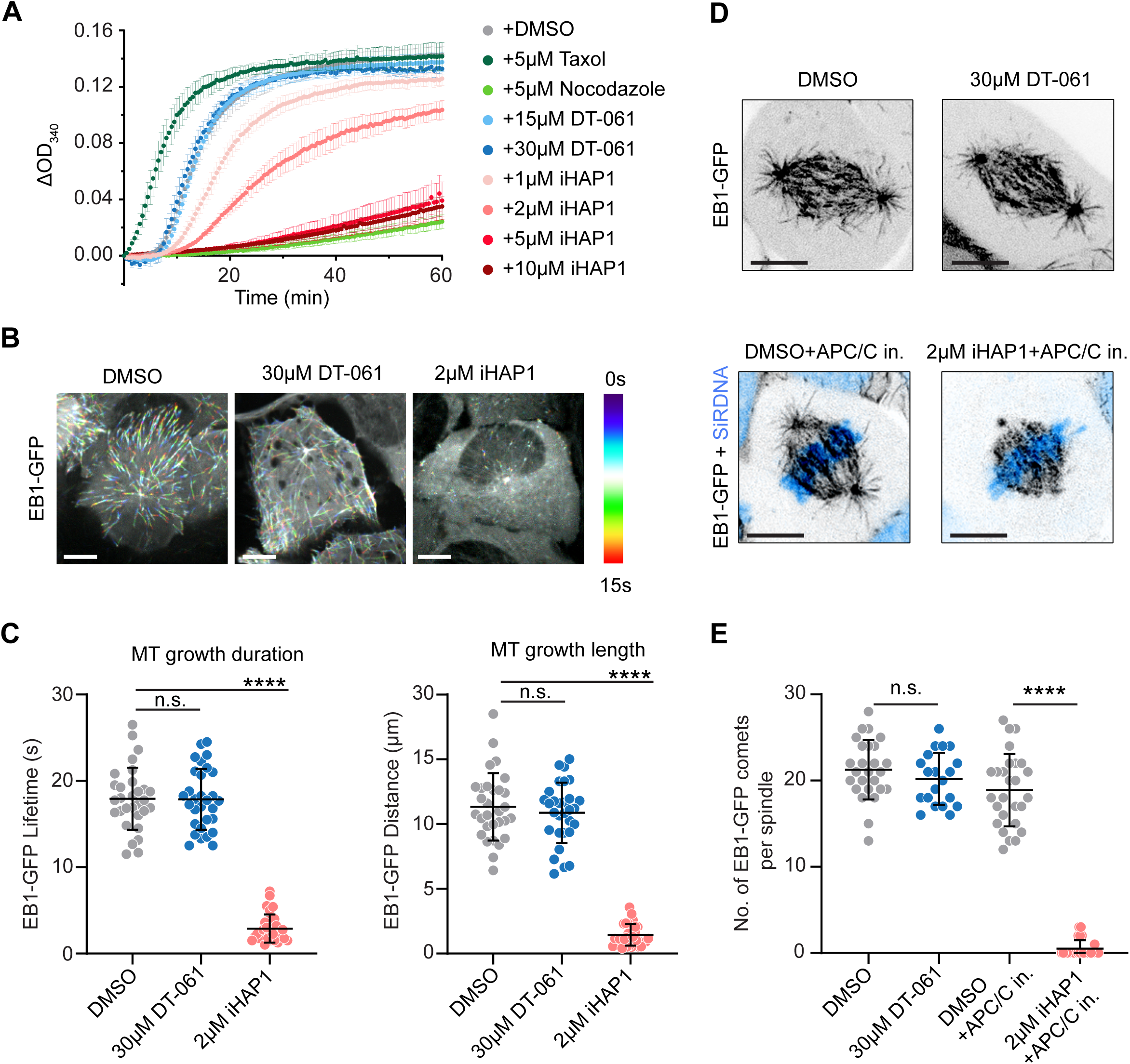
iHAP1 is a microtubule poison. A) *In vitro* tubulin polymerization assay showing concentration dependent effect of iHAP1 on polymerization dynamics. Taxol and Nocodazole served as controls for the experimental setup. The mean and SEM are shown from three to six independent experiments. B) Representative colour-coded temporal projection of U2OS EB1-GFP cells following indicated treatment. Scale bar 10µm. C) Quantification of microtubule dynamics acquired from manual tracking of EB1-GFP comets as indicators for MT growth. The mean and SD are plotted from three independent experiments (n = 10 cells per experiment). D) Representative temporal projection of mitotic spindles of U2OS EB1-GFP cells following specified treatment. DNA counterstained with SiR-DNA shown in cyan in merged images. Scale bar 10µm. E) Analysis of number of EB1-GFP comets per mitotic spindle. The mean and SD are plotted from three independent experiments (Total no. of cells = 26 (DMSO), 20 (30µM DT-061), 27 (DMSO+APC/C in.) and, 26 (2µM iHAP1+APC/C in).

To substantiate our *in vitro* results, we monitored the effect of 2 μM iHAP1 and 30 μM DT- 061 on microtubules in cells. We used a cell line expressing the EB1 microtubule tip tracking protein which allows direct monitoring of *in vivo* microtubule dynamics in a quantitative manner. The addition of iHAP1 to interphase cells for 20 minutes blocked microtubule growth while DT-061 had no effect (Fig. 3B-C, Movie 1). In agreement with its effect on microtubule dynamics, iHAP1 caused severe chromosome congression defects in mitotic cells shortly upon addition, with consequent spindle collapse and mitotic arrest (Fig. S7A, Movie 2-4). To prevent any potential effect of uncongressed chromosomes on analysis of microtubule dynamics in mitotic cells, we arrested cells in metaphase using APC/C inhibitors prior to iHAP1 treatment. Quantification of the number of EB1-GFP comets within astral microtubules revealed a similar strong effect of iHAP1 on microtubule growth in mitotic cells as well, where it nearly fully abolished astral microtubule polymerization (Fig. 3D-E). Astral microtubule dynamics in DT-061-treated mitotic cells remained unchanged. Moreover, we observed that iHAP1 led to a strong decrease in microtubule intensity on the mitotic spindle and absence of astral microtubules consistent with iHAP1 being a microtubule depolymerizing compound (Fig. S7B-D). Again, this effect on the mitotic spindle was observed 20 minutes after addition of iHAP1. Consistent with the rest of our data, DT-061 treatment did not cause any changes in spindle microtubule levels. Collectively, our *in vitro* and *in vivo* characterization uncovers iHAP1 as a microtubule poison, which is consistent with the mitotic arrest the compound induces. The observation that this happens *in vitro* and is nearly instantaneous in cells, argues that the effect of iHAP1 is independent of deregulated transcription.

We next analysed mitotic fidelity in iHAP1 treated cells and the impact of TACC3 and Rod1 removal. We used a HeLa cell line expressing fluorescent markers for tubulin and chromosomes and analysed mitotic progression by time-lapse microscopy (Fig. 4A-C). We used low doses of iHAP1 to monitor possible synergistic effects and recorded whether cells successfully progressed through mitosis or died during a mitotic arrest. Even though depletion of TACC3 and Rod1 causes quite distinct effects on mitotic timing, the proportion of cells dying during an iHAP1 induced mitotic arrest increased upon both TACC3 and Rod1 depletion (Fig 4A-C). This could explain the hypersensitivity to iHAP1 when TACC3 and Rod1 are depleted.

**Figure 4.**
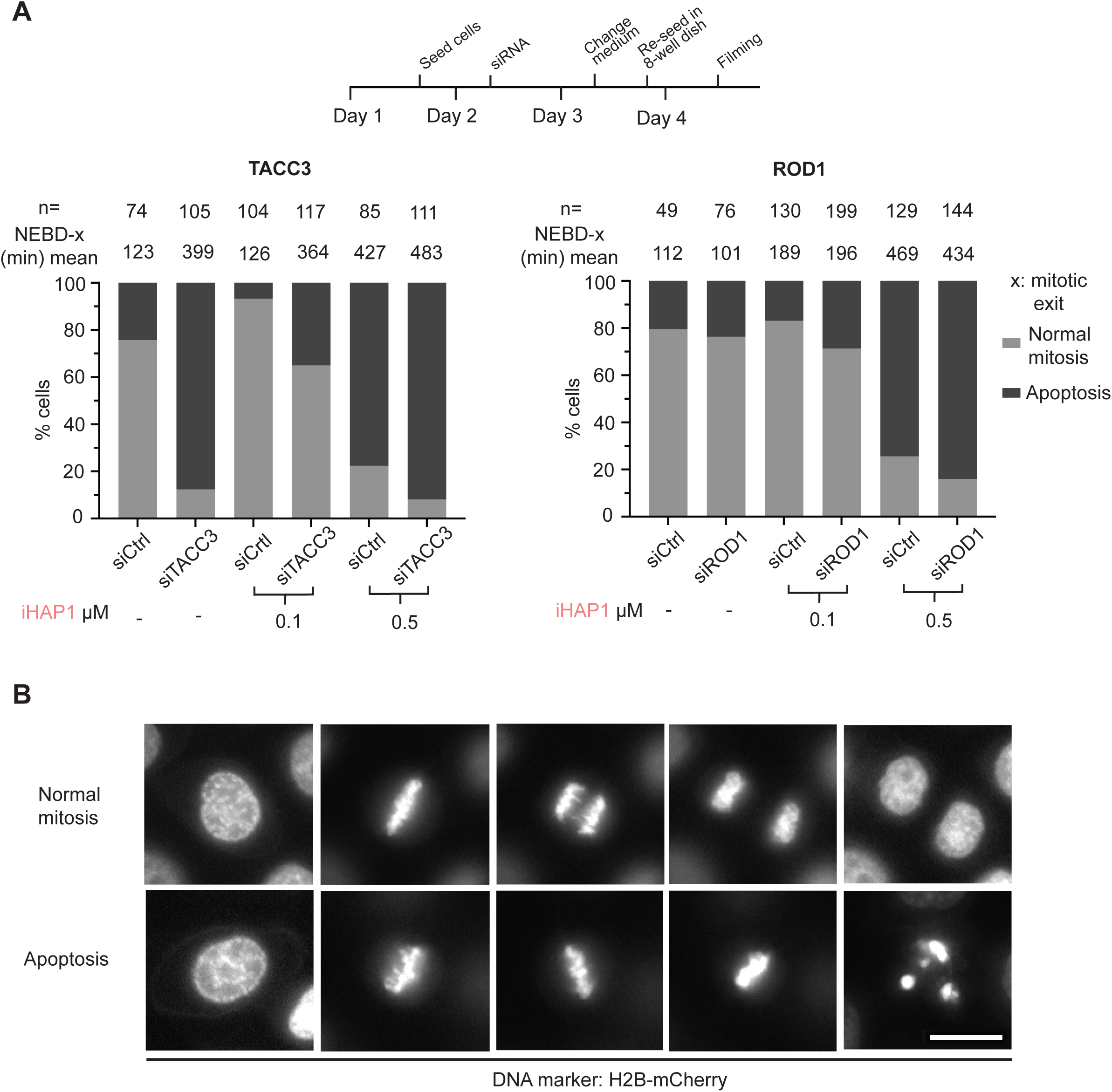
Depletion of TACC3 or Rod1 causes increased cell killing by iHAP1 during mitosis. A) Schematic of experimental protocol. B) Measurement of time spent in mitosis by time-lapse imaging taking into consideration whether cells exited mitosis or underwent cell death during mitosis. *n* indicates the number of cells analyzed from at least 2 independent experiments. C) Representative stills from time-lapse imaging experiments depicting the two phenotypes. Scale bar 20 µm.

In summary, all our data show that iHAP1 is a microtubule poison.

### DT-061 disrupts Golgi and ER

The TRAPP complex is known to regulate the transport between ER and Golgi while the NatC complex is required for N-terminal acetylation of proteins [30, 31]. Depletion of both TRAPP complex and NatC components has been shown to result in Golgi fragmentation [32–34]. Given that CRISPR-mediated removal of TRAPP and NAA complex components sensitized cells to DT-061, we investigated Golgi integrity in cells treated with DT-061. We used a live cell Golgi marker to investigate immediate effects upon DT-061 addition. Moreover, since many tricyclic compounds are fluorescent at low nm wavelength, which is also the case for DT-061 (Fig. S8A), we could directly visualize the sub-cellular localization of the compound. Strikingly upon addition of 20 μM DT-061 to HeLa cells, we observed a rapid disruption of the Golgi complex, which disintegrated into large vesicle-like structures that co-localised with DT-061 (Fig. 5A, Movie 5). We observed a similar effect on the Golgi complex in HeLa, RPE1, and MCF7 cells by immunostaining with an antibody against GM130, a marker for the cis-Golgi (Fig. S8B). Using a live cell marker for the ER, we observed that this diffuses to the nucleus shortly after addition of DT-061 (Fig. 5B, Movie 6). The effects of DT-061 on Golgi and ER markers were not observed with 2 μM iHAP1 (Fig. 5C-D). These results show that DT-061 affects both Golgi and ER integrity, consistently with the chemogenic profile of the compound. Considering the impact on Golgi and ER integrity, we analysed if DT-061 affects biosynthesis of sphingolipids as these biosynthesis pathways are tightly linked to these cellular structures (Fig. 6A)[35]. We fed breast carcinoma MCF7 cells with sphingosine, a precursor for sphingolipids, for four hours and then conducted quantitative mass spectrometry-based shotgun lipodomics to measure the induced increases in the levels of ceramide, sphingomyelin (SM), and glycosphingolipids hexosylceramide (isomers glucosyl- and galactrosylceramide) and ganglioside GM3. This was done in the presence of 15 μM DT-061, 1.5 μM iHAP1 and, as a control, we included Brefeldin A, which is known to inhibit vesicular transport between ER and Golgi, eventually causing the collapse of Golgi into ER [36]. In all our experiments, iHAP1-treated cells were similar to the control condition, indicating that iHAP1 does not affect the biosynthesis pathways of sphingolipids analysed here. For DT-061 and Brefeldin A, we observed a reduction in conversion of sphingosine to ceramide, a reaction occurring on the cytosolic side of the ER (Fig. 6B and Fig. S8C). Once transferred to the Golgi, ceramide can be modified into SM or glycospingolipids through addition of various headgroups. Conversion of ceramides to SM was increased by both DT-061 and Brefeldin A, while conversation to glycosphingolipids was specifically inhibited by DT-061. In contrast, we observed enhanced ceramide to glycosphinogolipid conversion in the presence of Brefeldin A. Interestingly, DT-061 and Brefeldin A had antagonistic effects on the viability of cells (Fig. S8D).

**Figure 5.**
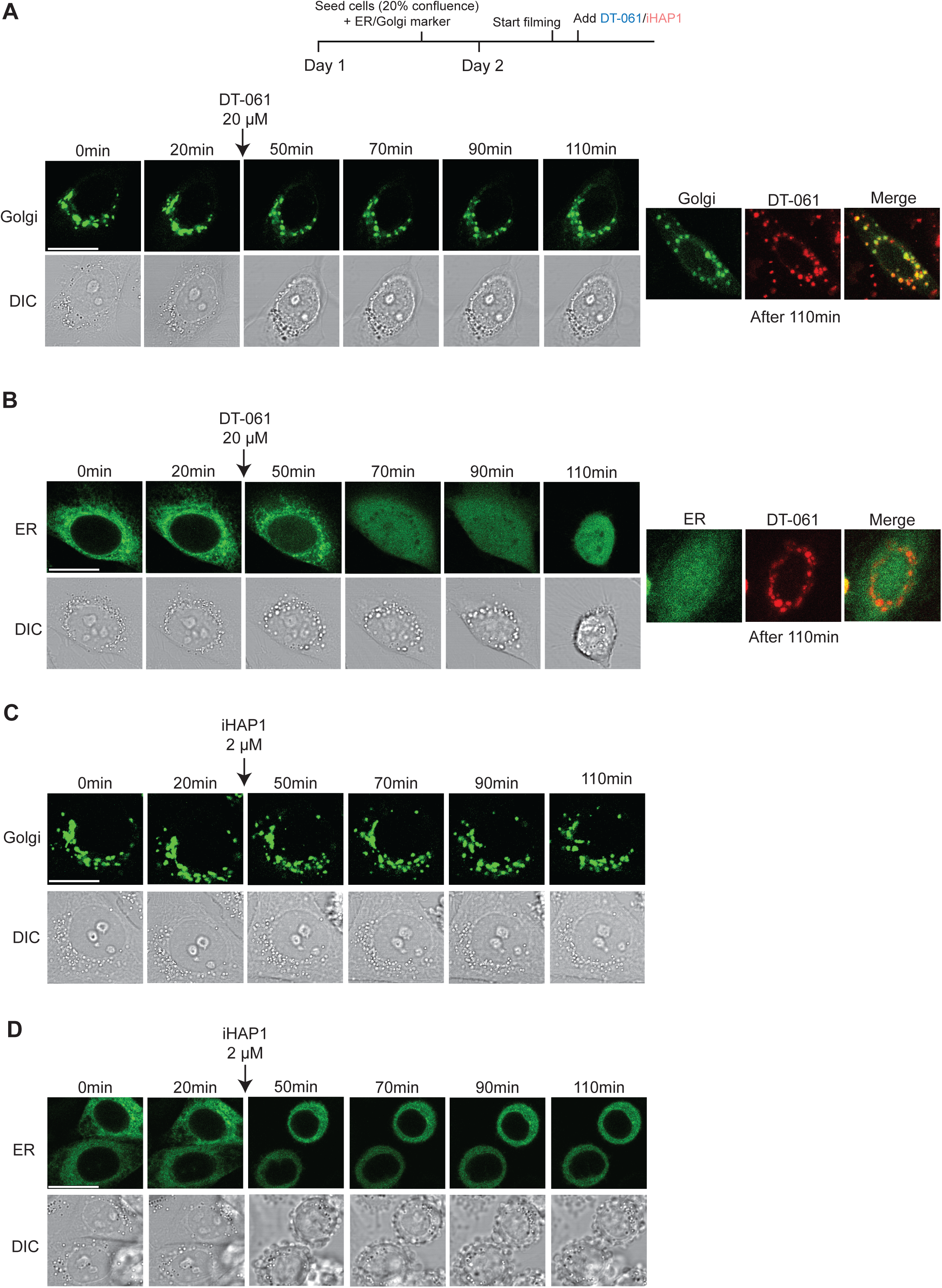
DT-061 disrupts the Golgi complex. A) Time-lapse imaging of HeLa cells expressing a marker staining the Golgi apparatus (green). After 20 minutes, DT-061 was added. On the right-hand side, the co-localisation of DT-061 (red) and the Golgi marker is shown. B) As in A) but with a marker for ER. C-D) as A) but iHAP1 added. Representative stills from at least 2 independent experiments. Scale bar 20 µm.

**Figure 6.**
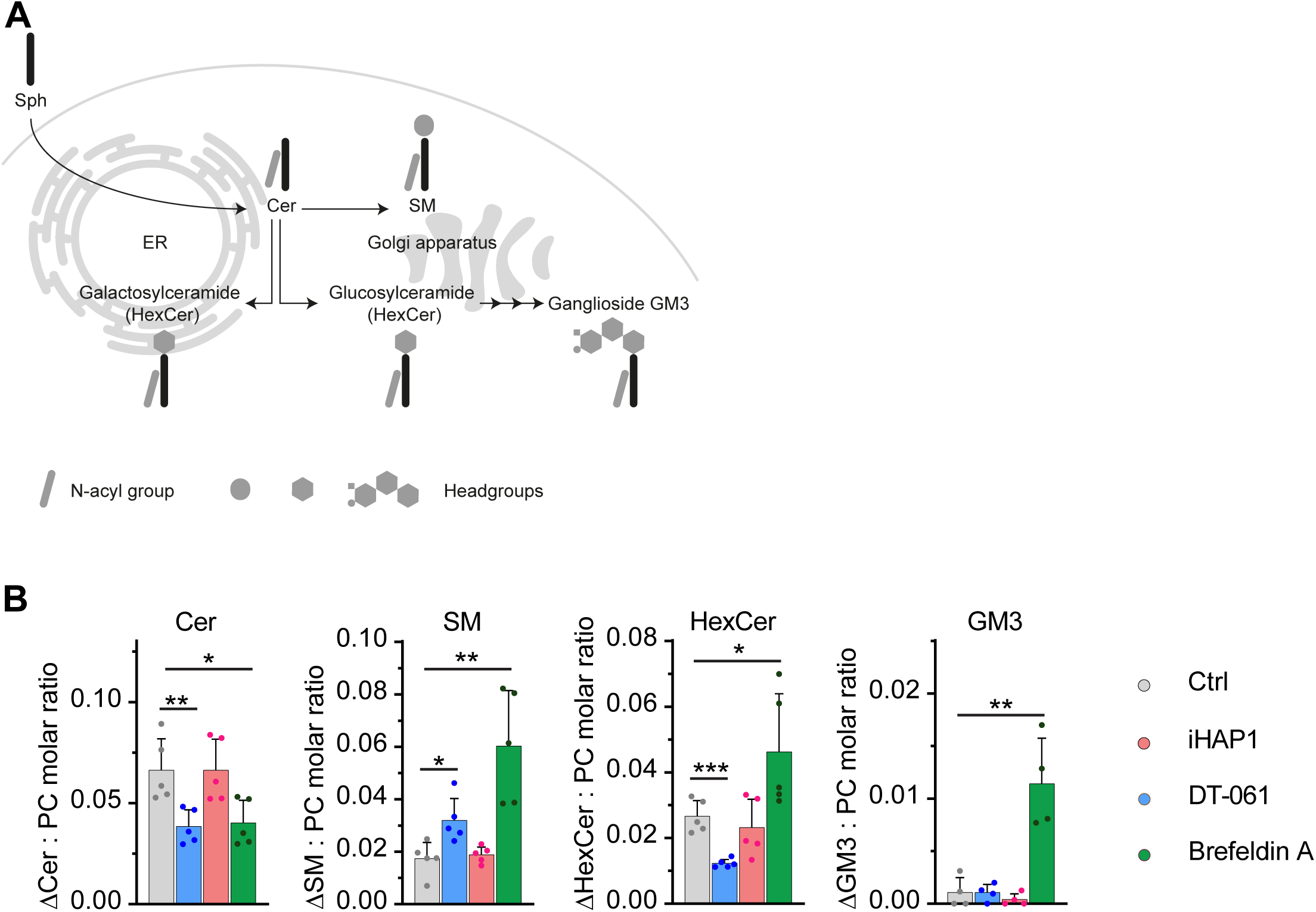
DT-061 affects sphingolipid biosynthesis in Golgi and ER. A) Schematic illustration of cellular sphingolipid biosynthesis in ER and Golgi. B) Elevations in the molar quantities of sphingolipid classes detected in MCF7 cells after feeding with the precursor sphingosine. MCF7 cells were co-treated with DT-061, iHAP1, Brefeldin A, or vehicle. The determined molar quantities of sphingolipid classes were normalized to that of PC. The presented values represent averages of five independent experiments. *p<0.05, **p<0.01, ***p<0.001. The abbreviations used are Cer: ceramide, HexCer: hexosylceramide, PC: phosphatidylcholine, and SM: sphingomyelin.

Collectively, our work shows that DT-061 has a clear and immediate effect on Golgi and ER integrity which impacts lipid biosynthesis pathways occurring at these cellular structures.

## Discussion

PP2A dampens the activity of oncogenic signaling pathways and for this reason there is a considerable interest in strategies able to activate PP2A holoenzymes. Here, we extensively analysed two tricyclic heterocyclic compounds that have been claimed to activate specific PP2A-B56 complexes to inhibit cellular proliferation [10, 11]. Collectively, our work shows that none of these compounds have a specific effect on PP2A-B56 and that their antiproliferative effects are unrelated to PP2A-B56 activation.

The molecular targets of iHAP1 are microtubules, and this compound has a clear effect on microtubule stability both *in vitro* and *in vivo*. This has already been documented by a previous study characterizing iHAP1 and related compounds [29]. Why this effect was not observed by Morita et al. (2020) in their *in vitro* experiments, is unclear to us. The fact that iHAP1 is a microtubule poison is consistent with the cellular phenotypes observed in the present work and by Morita et al. (2020), namely a strong mitotic arrest. However, we disagree on the molecular mechanism behind the mitotic arrest. Morita et al. (2020) argued that the effect is mediated through PP2A-B56ε dephosphorylation of MYBL2, which results in a deregulation of mitotic proteins. We see an immediate effect of iHAP1 on microtubule dynamics in interphase and mitotic cells arguing against any transcriptional effects. We note that no direct binding between iHAP1 and PP2A-B56ε was shown by Morita et al. (2020).

The effect of DT-061 on Golgi and ER integrity is striking and fully consistent with the chemogenetic profile of the compound established by our CRISPR screen. Depletion of TRAPP and NatC components is known to affect Golgi integrity, which in turn sensitizes these cells to further perturbation of the Golgi and ER by DT-061. At present, we do not know the direct molecular target of DT-061, but we favor the idea that it is a lipid constituting part of the Golgi or ER. This would be consistent with the hydrophobic nature of DT-061 and the fact that we see DT-061 forming small structures in cells attracting the Golgi marker. Tricyclic heterocyclic compounds can have a profound impact on lipids and indeed methylene blue, which belongs to this class of molecules, has been used to stain the Golgi in cells [37–41]. We acknowledge the extensive studies conducted by Leonard et al. (2020) to analyse the interaction between DT-061 and PP2A-B56α complexes. Our analysis of the published cryo-EM structure of DT-061 bound to PP2A-B56α argues that the flexible C-terminal tail of the catalytic subunit could also be modelled into the density ascribed to DT-061. It is important to point out that the cryo-EM structure is the most compelling evidence for direct binding of DT-061 to PP2A-B56α. The biochemical experiments in Leonard et al. (2020) focused on the stabilization of PP2A complexes by DT-061 which could be unrelated to direct binding. We were unable to recapitulate a stabilizing effect on the PP2A-B56α holoenzyme as observed by Leonard et al. (2020) either *in vitro* or *in vivo*. Our extensive analysis of DT-061 reveals that its cellular toxicity is related to disruption of Golgi and ER function.

Collectively, our work argues that the PP2A activating properties of iHAP1 and DT-061 and possibly other tricyclic compounds should be carefully reconsidered. Previous literature using such compounds to make claims on PP2A regulation should also be revisited.

## Acknowledgements

Work at the Novo Nordisk Foundation Center for Protein Research is supported by grant NNF14CC0001. JN is supported by grants from the Danish Cancer Society (R269-A15586-B17), Independent Research Fund Denmark (8021-00101B and 0134-00199B) and Novo Nordisk Foundation (NNF18OC0053124 and NNF20OC0065098). The work was supported by Independent Research Fund Denmark (grant no. 7016-00055) to NM and 6108–00542B to KM. JD is supported by the Novo Nordisk Foundation (grant agreement no. NNF18CC0033876). MW was supported by the Swiss National Fund (P2EZP3_178624) and the Danish Lundbeckfonden (2017-3212). Work in the lab of MB is supported by grants from the Danish Cancer Society (R146-A9322), the Lundbeck Foundation (R215-2015-4081) and the Novo Nordisk Foundation (NNF19OC0058504). This work is supported by grants R35GM119455 from NIH/NIGMS and R33CA225458 from NIH/NCI to ANK. The Novo Nordisk Foundation Center for Stem Cell Biology is supported by a Novo Nordisk Foundation grant number NNF17CC0027852. We thank Martina Barisic for technical assistance. Porcine brain for tubulin purification was kindly provided by Karsten P Hammelev, Copenhagen University, Denmark. NMR data was recorded at cOpenNMR, an infrastructure facility funded by the Novo Nordisk Foundation (#NNF18OC0032996). We thank the NNF CPR protein production facility for help with expression and purification of PPP2R1A. We thank Magali Michaut and the DanStem Genomics Platform for technical expertise, support, and the use of instruments. Data processing and analysis were performed using the DeiC National Life Science Supercomputer at DTU (www.computerome.dk). We thank Iain Cheeseman for providing the inducible CRISPR Cas9 cell lines. We thank Goutham Narla for providing DT-061. We thank Jukka Westermarck, Maja Koehn and Veerle Janssens for fruitful discussions.

## Author contributions

GV performed *in vitro* and *in vivo* analysis of compounds and CRISPR screen validation. JD performed imaging experiments of mitosis with depletions and imaging of Golgi and ER. GR performed all experiments in Fig. 3 and Fig. S7. ETH performed the CRISPR screen with the help of GV and JN. BCM and ANK performed mass spectrometry analysis and microcystin-bead experiments. KM and LKKH performed lipidomics experiments. MBW, GM analysed the cryo-EM structure. BLM performed ITC experiments and stability analysis of PPP2R1A. JN, MB and NM coordinated the work in their labs. JN coordinated the overall work and wrote the manuscript.

## Methods

### iHAP1 and DT-061

We obtained iHAP1 from Enamine (Cat#: Z56843374) and DT-061 (Cat#: HY-112929) from either MedChemExpress or as a gift from Goutham Narla. Compounds were dissolved according to instructions and stored as single use aliquots. Their identity was confirmed by mass spectrometry.

### Antibodies

Following antibodies were used at the indicated dilutions: Myc tag (#ab32, 1:1000, Abcam), PP2A scaffold subunit (#2041, 1:2500, Cell Signaling Technology), PPP2R2A (#5689, 1:2500, Cell Signaling Technology), PPP2R5A (#610615, 1:3000, BD Biosciences), PPP2R5D (#5687, 1:2500, Cell Signaling Technology), PPP2R5E (#PA5-17981, 1:2000, Thermo Fisher Scientific), PP2AC Methyl (Leu 309) (#828801, 1:3000, BioLegend), actin, (#sc-47778 HRP, 1:5000, Santa Cruz Biotechnology), TRAPPC2L (#HPA041714, 1:2000, Sigma-aldrich), TACC3 (sc-376883, 1:1000, Santa Cruz Biotechnology), ROD1 (1:300, a kind gift from Reto Gassmann’s lab), GM-130 (#610822, 1:500, BD Biosciences), Alexa Fluor 488 (#A32723, 1:1000, Thermo Fisher Scientific), Tubulin (#T9026, DM1A, 1:2000, Sigma-Aldrich).

### Cell culture methods

Human cell cultures were maintained at 37°C in a humidified 5% CO_2_ environment. Cells were cultured in the appropriate medium supplemented with the necessary nutrients as indicated by the American Type Culture Collection, Manassas, Virginia, U.S.A. Cell media were supplemented with 10% Fetal Bovine Serum (FBS) (Thermo Fisher Scientific) and 1% Penicillin-Streptomycin 10,000 U/ml, (Thermo Fisher Scientific). Media used for RNAi experiments was devoid of Penicillin-Streptomycin.

CRISPR/Cas9-inducible knockout cell lines were cultured in presence of tetracycline-free FBS (Thermo Fisher Scientific). To activate Cas9 activity, cells were induced with Doxycycline at a concentration of 1 µg/mL for 4 days. Downregulation/knock-out of the protein of interest was assessed by Western blotting.

Parental U2OS and U2OS cell lines stably expressing H2B-GFP and mCherry-α tubulin (gift from S. Geley, Innsbruck Medical University, Austria)[42], EB1-GFP [43] (gift from P. Draber, IMG ASCR, Prague, Czech Republic) were grown in Dulbecco’s Modified Eagle Medium (DMEM; Gibco) supplemented with 10% fetal bovine serum (FBS; Invitrogen) at 37°C with 5% CO_2_.

U2OS EB1-GFP cells were treated with DT-061 and iHAP1 20 minutes before imaging. To arrest cells in metaphase, cells were incubated with the Anaphase-Promoting Complex/Cyclosome (APC/C) inhibitors Apcin (20µM) and a cell permeable tosyl-L-arginine methyl ester (proTAME) (10µM) for 2 hours before the addition of DT-061 or iHAP1. DNA was labelled by adding 20 nM SiR-DNA (Spirochrome) 2 hours prior to live-cell imaging.

### *In vitro* phosphatase assay

PP2A holoenzymes were purified from HeLa cell lines stable expressing FLAG-tagged B55α, B56α and B56ε regulatory subunits, respectively, which were previously generated in the lab[6, 44]. Cells were grown in 15 cm dishes in presence of 10 ng/mL Doxycycline for 48 hours and washed 3 times in 50 mL ice-cold PBS. A total of ∼ 8×10^7^ cells were lysed in 2 mL lysis buffer (20 mM Tris HCl pH 7.4, 150 mM NaCl, 1% Triton-X 100 supplemented with protease inhibitor Complete, Roche). Lysates were rotated at 4°C for 20 minutes and subsequently centrifuged at maximum speed for 30 minutes. Anti-FLAG M2 Affinity gel (Sigma-Merck) was equilibrated in lysis buffer and gently shaked at 4°C for 3 hours with the lysate. Beads were subsequently washed 2 times in lysis buffer and 4 times in TBS/0.1 % Triton-X 100, followed by one wash in TBS. Prior to elution, beads were washed one time in elution buffer (20 mM Tris-HCl pH 7.4, 150 mM NaCl, 1 mM MnCl_2_). Elution of the co-immunoprecipitated complex was performed by gentle shake of the beads at 4°C in elution buffer supplemented with 3X FLAG peptide (Sigma-Aldrich) for 30 minutes. Glycerol and DTT were added at a 25% and 1 mM final concentration, respectively, before storing the samples at -80°C. Non-stick tubes were used during the purification and for storage. Purity of enzymatic preparations was assessed by colloidal staining (NOVEX Colloidal Blue Stain Kit, Invitrogen) and Mass Spectrometry. No other protein phosphatases were detected in the preparations. Titration curves were performed using different amounts of enzyme (2-10 µL).

*In vitro* reactions were performed in non-stick tubes. 2 µL of enzyme were incubated with 1 mM basic (WRRA(pT)VA) or acidic (WSGDD(pT)IVD) peptide substrate together with 20 µM DT-061 or 10 µM iHAP1 in phosphatase buffer (50 µM Tris-HCl pH 7.4, 0.3 mM NaCl and 2 mM MnCl_2_) for 15 minutes at 30°C. Peptide were purchased from Peptide 2.0 Inc (Chantilly, VA, USA). Inorganic phosphate release was assessed using the PiColorLock Phosphate Detection System (Expedon) according to manufacturer’s instructions. Data from three independent experiments were analysed.

### Protein production and purification

GST-PPP2R1A was expressed in *E. coli* and cells lysed in buffer L (50 mM Tris-HCl pH=7,5; 300 mM NaCl, 10% glycerol, 0.5 mM TCEP). Following lysis and clarification of the extract, it was loaded on a glutathione column and washed with buffer L. The fusion protein was eluted with buffer L containing 20 mM reduced glutathione and GST tag removed by TEV cleavage. The GST was removed by loading the TEV cleavage reaction on a glutathione column. Untagged PPP2R1A was further purified on a Superdex 200 16/60 equilibrated with buffer G (50 mM NaP pH=7.5, 150 mM NaCl, 5% glycerol, 0.5 mM TCEP) and peak fractions were pooled. The protein was analysed by LC-MS.

PP2A-B56α was reconstituted in HEK293 cells by co-expressing Strep-B56α, His-PP2ACA and MBP-PPP2R1A. The cell pellet was lysed in 100 mM Tris pH=8.0, 150 mM NaCl, 10% glycerol, 0.5 mM TCEP and the complex purified by a Strep-tag affinity column and the peak fractions were run on GF200 10/30 column and collected and frozen.

### NMR

1D-^1^H-NMR spectra with water suppression (bruker-pulseprogram: zgesgp) were recorded at 25 °C on a Bruker Avance III HD spectrometer equipped with a 5mm TCI Cryoprobe H-C/N-D. The final protein concentration was 20 µM in all samples and the compounds were added to a final concentration of 200 µM from a d_6_-DMSO stock. The buffer composition in all samples were 116 mM KCl, 2.3 mM Na_2_HPO_4_, 1.7 mM KH_2_PO_4_, 10% d_6_-DMSO and 5% D_2_O.

### Isothermal titration calorimetry (ITC)

Prior to ITC experiments the recombinant PPP2R1A protein was extensively dialyzed against ITC buffer, 100 mM sodium phosphate pH 7.5, 150 mM NaCl, 0.5 mM TCEP. After dialysis, the ITC buffer was supplemented with either 5% or 10% DMSO (Sigma-Aldrich, (#34943)) before running the ITC experiments. Both DT-061 and iHAP were dissolved in the same dialysis buffer from an initial stock 1 mM in pure DMSO to a final concentration of 20 μM and adjusting the percentage of DMSO to 5% and 10% with pure DMSO. All ITC experiments were performed on an Auto-iTC200 microcalorimeter (MicroCal-Malvern Panalytical, Malvern, UK) at 25 °C. The protein was loaded into the syringe at 300 μM and titrated into the calorimetric cell containing the DT-061 or iHAP1 solutions at 20 μM. Control experiments with the protein injected in the sample cell filled with buffer were carried out under the same experimental conditions. In all assays, the titration sequence consisted of a single 0.4 μl injection followed by 19 injections, 2 μl each, with 150 s spacing between injections to ensure that the thermal power returns to the baseline before the next injection. The stirring speed was 750 rpm and the reference power set at 8 μcal/s. The heats evolved after each ligand injection were obtained from the integral of the calorimetric signal.

### Differential scanning fluorimetry (nanoDSF)

The thermal stability of PPP2R1A in both NMR and ITC buffers with different percentage of DMSO (from 0% to 40%) was evaluated by following the changes in the intrinsic fluorescence ratio (350/330 nm) as a function of the temperature. The experiments were run in a Prometheus NT.48 nanoDSF (NanoTemper Technologies). 10 μl of the protein samples at 10 μM were measured in triplicates with a temperature ramp set to 1 °C/min from 20 °C-80 °C using standard treated capillaries (NanoTemper Technologies (#PR-C002)). The PR.ThermControl v2.1.6 software (NanoTemper Technologies) was used for data acquisition and data analysis.

### Mass photometry (MP)

Mass photometry experiments were performed on a Refeyn One^MP^ (Refeyn Ltd) mass photometer at room temperature. Measurements were run in triplicates diluting the protein sample at 50 nM concentration in NMR buffer loaded on a gasket (Grace Bio-Labs reusable CultureWell^TM^ gaskets, Merck (#GBL103250)) mounted on a glass microscope coverslip (Marienfeld (#0107222)). The molecular weight was obtained by contrast comparison with known molecular weight mass standard calibrants (NativeMark^TM^ Unstained Protein Standard, ThemorFischer Scientific, (#LC0725)) measured on the same buffer and on the same day. Movies were recorded using AcquireMP (Refeyn Ltd, version 2.3.2), with a regular field of view and exposure time of 0.95 ms for the acquisition camera settings. A total of 6000 frames were recorded over a duration of one minute. Movies were processed and analysed with the DiscoverMP software (Refeyn Ltd, version 2.3.0) provided by the instrument manufacturer.

### Immunoprecipitation

HEK-293T cells stably expressing inducible myc-PP2AC subunits [45] were induced with 10 ng/mL Doxycycline 48 hours before immunoprecipitation. After treatment with DT-061 20 µM, iHAP1 2µM or DMSO for 30 minutes, cells were lysed in low salt lysis buffer (50 mM NaCl, 50 mM Tris pH 7.4, 1 mM EDTA, 0.1% NP-40, 1 mM DTT) supplemented with Complete protease inhibitor (Roche). Lysates were rotated at 4°C for 20 minutes prior to maximum speed centrifugation at 4°C. Clear protein lysate was transferred to a new, non-stick prechilled reaction tube to perform immunoprecipitation. Myc-Trap Agarose affinity reagent (Chromotek) and Anti-FLAG M2 Affinity gel (Sigma-Merck) were used to immunoprecipitate PP2A catalytic subunit and PP2A regulatory subunits B55α, B56α and B56ε, respectively. Beads were equilibrated in lysis buffer and then rotated for 1 hour in the protein lysate of interest at 4°C. Beads were subsequently washed 3 times in lysis buffer and bound complexes were eluted using 4x Lämmli sample buffer (Biorad). Samples were boiled at 95°C for 7 minutes and resolved through SDS-PAGE prior to Western blotting or analysed by quantitative Mass Spectrometry as described in the label-free LC-MS/MS analysis section. Experiments were performed in triplicate.

### Label-free LC-MS/MS analysis

**PIB pull-downs** were analysed on a Q-Exactive Plus quadrupole Orbitrap mass spectrometer (ThermoScientific) equipped with an Easy-nLC 1000 (ThermoScientific) and nanospray source (ThermoScientific) as previously described [15]. Raw data were searched using COMET (release version 2014.01) in high resolution mode [46] against a target-decoy (reversed) [47] version of the human proteome sequence database (UniProt; downloaded 2/2020) with a precursor mass tolerance of +/- 1 Da and a fragment ion mass tolerance of 0.02 Da, and requiring fully tryptic peptides (K, R; not preceding P) with up to three mis-cleavages. Static modifications included carbamidomethylcysteine and variable modifications included: oxidized methionine. Searches were filtered to a <1% FDR at the peptide level. Quantification of LC-MS/MS spectra was performed using MassChroQ [48] and the iBAQ method [49]. PPP subunit abundances were normalized across all samples by quantile normalization. Statistical analysis was carried out in R statistical software (version 4.0.5) in conjunction with R studio (version 1.3) (Team, 2019) by two-tailed Student’s t-test.

**B56a, B56e, and B55a** purifications were analysed on an Orbitrap Fusion Lumos mass spectrometer (ThermoScientific) equipped with an Easy-nLC 1200 (ThermoScientific), searched, filtered, and quantified as described above.

**PP2Ac pulldowns** were analysed on an Orbitrap Fusion Lumos mass spectrometer (ThermoScientific) equipped with an Easy-nLC 1200 (ThermoScientific), searched, filtered, and quantified as described above. PP2Ac abundance was normalized across all samples to be equal. Statistical analysis was carried out in R statistical software (version 4.0.5) in conjunction with R studio (version 1.3) (Team, 2019) by two-tailed Student’s t-test.

### Analysis of Cryo-EM map

Cryp-EM maps and PDBS were visualized with Chimera [50], superpositions were done in pymol (The PyMOLMolecular Graphics System, version 1.2r3pre, Schrödinger, LLC) and the C-terminal tail of PP2AC was built in Coot [51] and refined in phenix [52].

### CRISPR-Cas9 knockout screens

Viral particles of the LCV2:TKOv3 sgRNA library were produced as previously described [53]. RPE1-hTERT Flag-Cas9 TP53-/- cells were transduced with an MOI of 0.3 at a coverage of > 350-fold sgRNA representation, which was maintained throughout the screen at each cell passage point. Twenty-four hours after transduction, transduced cells were selected for 24 hours with 25 µg/ml puromycin. Cells were treated with trypsine and reseeded in the same plates while maintaining puromycin selection for another 24 hours. Three days after transduction, which was considered the initial time point (T0), cells were pooled and passaged while cell pellets of two replicates of 3.5×10^7 cells were frozen for downstream processing. Cells were passaged after three and nine days after transduction, respectively, which was considered T6, cells were split into technical duplicates of either mock treated conditions or treated with DT-061 (5.5 µM) or iHAP1 (0.5 µM) equivalent to pre-determined LD20 concentrations in uninfected RPE1-hTERT Flag-Cas9 TP53-/- cells treated for 12 days. Cells were subcultured every three days (T9, T12 and T15) in medium with or without drugs until the final time point at T18 at which point cell pellets from 3.5×10^7 cells were frozen from each replicate. Genomic DNA from cells collected at T0 and T18 was isolated as previously described [54] and sgRNA sequences amplified by PCR using Q5 Mastermix Next Ultra II (New England Biolabs, Cat# M5044L) with the following primers: LCV2_forward: 5’- GAGGGCCTATTTCCCATGATTC -3’ and LCV2_reverse: 5’- GTTGCGAAAAAGAACGTTCACGG -3’. This was followed by a second PCR reaction containing i5 and i7 multiplexing barcodes and final gel-purified products were sequenced on Illumina NextSeq500. Fastq files were generated using bcl2fastq v2.19.1 and reads were trimmed to 20 bp using cutadapt 1.18 [55] removing a variable number of bp at start and end depending on the size of the primer stagger. MAGeCK 0.5.8 [56] was used to assign the trimmed reads to the guides in the TKOv3 library and create the count matrix. Gene scores (normZ values) were estimated from the count matrix using the drugZ algorithm [25]. To assess data quality, we generated precision-recall curves, calculated by the BAGEL.py ‘pr’ function [57] using the core essential (CEGv2.txt) and nonessential (NEGv1.txt) gene lists from (https://github.com/hart-lab/bagel), comparing T0 to T18 for mock treated cells.

### Tubulin polymerization assay

Assembly competent tubulin was purified from porcine brain as described before [58]. Turbidity based microtubule polymerization assay was performed as described before [59]. Briefly, indicated concentrations of iHAP1 and DT-061 or DMSO (vehicle control) were mixed with free tubulin (2mg/ml final concentration) and microtubule assembly was induced by the addition of 1mM GTP and 10% glycerol in BRB80 buffer (80mM PIPES, pH 6.8, 1mM MgCl_2_, and 1mM EGTA) at 37°C. Paclitaxel (taxol), a polymerization enhancer and nocodazole, a microtubule depolymerizer were used as internal controls for the polymerization reaction. Microtubule polymerization was monitored by measuring the change in absorbance (340nm) using SpectraMax® iD3 (Molecular Devices) microplate reader in 30 seconds intervals over 60 minutes.

### Live-cell imaging

Time-lapse imaging was performed as described before [60]. Briefly, cells were cultured in 35 mm glass-bottomed dishes (14 mm, No. 1.5, MatTek Corporation) and imaging was performed in a environment controlled chamber (37 °C with controlled humidity and 5% CO_2_ supply), using a Plan-Apochromat DIC 63x/1.4NA oil objective mounted on an inverted Zeiss Axio Observer Z1 microscope (Marianas Imaging Workstation from Intelligent Imaging and Innovations Inc. (3i), Denver, CO, USA), equipped with a CSU-X1 spinning-disk confocal head (Yokogawa Corporation of America) and four laser lines (405 nm, 488 nm, 561 nm and 640 nm). Image detection was performed using an iXon Ultra 888 EM-CCD camera (Andor Technology).

Mitosis in U2OS-H2B-GFP/mCherry-α tubulin cells were imaged using fifteen 1 μm-separated z-planes collected every 2 minutes. Microtubule dynamics were monitored by imaging U2OS EB1-GFP cells every 500ms. 2-5 EB1 comets per cell were tracked manually using ImageJ. In both experiments, iHAP1 or DT-061 were added 20 minutes before imaging.

### Immunofluorescence staining

Cells were treated with siRNA against TRAPPC2L 72 hours before microscopy as explained in the corresponding methods section. Cells were seeded on coverslips 24 hours before immunofluorescence staining. Cells were then fixed in 4% paraformaldehyde (PFA) (Thermo Fisher Scientific) solution for 10 minutes and subsequently permeabilized with 0.2% triton X- 100 (Sigma-Aldrich) for 5 min, washed once in PBS and blocked for 1 hour in 3% Albumin (Sigma-Aldrich) in PBS. Subsequently, cells were incubated with an antibody against GM-130 (see antibody section) for 1 hour. Three washes with PBS followed, each for 5 minutes. Species-specific secondary antibody was then incubated for 30 min in the dark. DNA was counterstained with DAPI solution (#62248, 1:2000, Thermo Fisher Scientific) diluted in PBS, together with the secondary antibody (see antibody section). The coverslips were subsequently washed 3 times with PBS and briefly rinsed with 100% reagent grade ethanol. Coverslips were then mounted in Vectashield antifade mounting medium (Vector Laboratories) and were analyzed by fluorescence microscopy. Stills with z-stacks 200 nm apart were taken using the DeltaVision Elite system microscope (GE Healthcare), 100x oil immersion objective, 1×1 bin. The same settings were applied for all immunostaining stills: DAPI 5% laser power (0,025 exposure), FITC 10% laser power (0,2 exposure), followed by deconvolution using Softworx. U2OS cells were treated with the APC/C inhibitors Apcin (20µM) and proTAME (10µM) for 2 hours followed by DT-061 or iHAP1 for 20 minutes, fixed and stained as described before [61]. Mouse anti α-tubulin was used as primary antibody (see antibody section).

### Transient siRNA transfection

Silencer Select Pre-designed siRNA oligomers were purchased from Life Technologies LTD. Following antisense oligos were transfected as described below: TRAPPC2L (#s28534), ROD1 (#s18778), TACC3 (#s20471). A non-targeting oligo against Luciferase was used as control. The oligos were briefly centrifuged and resuspended in nuclease-free water according to the manufactureŕs instructions. Seeding of cells was performed at the same time of transfection. 200 μl Opti-MEM medium (Gibco) was mixed with 2 μl of a 1 nMol stock of the oligonucleotides and briefly resuspended. 4 μl of Lipofectamine RNAiMAX Transfection Reagent (Thermo Fisher Scientific) was added to the solution and incubated at room temperature for 15 minutes. Subsequently, the mixture was added dropwise on the fresh-seeded cells in a ∼10 cm^2^ culture dish area in 1.8 mL Opti-MEM medium and incubated for 72 hours.

### Transient siRNA transfection and live imaging

HeLa stabling expressing Tubulin-mVenus, H2B-mCherry (gift from Barr Lab, Institute of Clinical Sciences, Imperial College, London) were transiently transfected with the following antisense oligos: ROD1 (5′ GGAAUGAUAUUGAGCUGCUAACAAA 3′; ThermoFisher Scientific), TACC3 (5’ GAGCGGACCUGUAAAACUA 3’) and a non-targeting oligo against Luciferase was used as control (Sigma, Custom order). RNAi Max (Invitrogen) was used according to manufactures instructions.

Live cell imaging of cells depleted of TAAC3 and ROD1 was performed using a DeltaVisionElite system fluorescent microscope (GE Healthcare) equipped with a CoolSNAP HQ2 camera (Photometrics), 40x oil immersion objective.

Cells were seeded in an 8-well ibidi dish (Ibidi) the day before filming, the media was changed to Leibovitz’s L-15 (Life Technologies) immediately before the filming. YFP (10% laser power, exposure 0,2) and mCherry (5% laser power, 0,05 exposure) were recorded at 7 minutes intervals and data were analyzed using SoftWoRx (GE Healthcare). The time from nuclear envelope breakdown (NEBD) to anaphase was measured in single cells from independent experiments. Further analysis carried out with GraphPad Prism 9 (GraphPad Software, San Diego, California, USA).

### Cell growth assays

The Sulforhodamine B (SRB) proliferation assay was used to assess *in vitro* cytotoxicity of the compounds [62]. Cells were treated with sublethal drug concentrations (LD20) for 12 days from the day of seeding. If necessary, transient siRNA transfection was performed three days before drug treatment. After 12 days, medium was removed from the wells and cells were washed once with PBS and fixed in prechilled 10% trichloroacetic acid (TCA) (Sigma-Aldrich) for 30 minutes at 4°C. Cells were subsequently washed two times with double-distilled water (dd-H_2_O) and stained with 0.4% SRB (/1% acetic acid for 20 minutes at room temperature protected from light. SRB was then removed, and cells were washed four times in 1% acetic acid. After complete light-protected drying, SRB was dissolved by addition of 10 mM Tris pH 8 and gentle shaking at room temperature for 2 hours. 100 µL of the solution were then transferred to a 96-well plate and absorbance at A_510_ was read with plate reader. Percentage of cell-growth inhibition was calculated in relation to a mock-treated control. The IncuCyte^®^ live-cell analysis system was additionally used to real-time track the growth of the cells subjected to the different drug treatments and gene knock-down.

### Live imaging of DT-061-treated cells

HeLa parental cells were seeded in 8-well dish (Ibidi) and incubated for a minimum of 16hours with the fusion construct CellLight™ Golgi-GFP or CellLight™ ER-RFP, BacMam 2.0 (Invitrogen™). Right before imaging, media was exchanged to Leibovitz’s L-15 (Life Technologies). DT-061 (MCE MedChem Express) was added to the ongoing experiment for a final concentration of 20-40µM.

Filming was performed on a LSM 880 Airyscan Confocal attached to an inverted stand Zeiss AxioObserver.Z1 (Carl Zeiss), equipped with an Incubation box XL Multi S1 set to 37°C, and sample was mounted on a C-Apochromat x40/1.2W objective. Time-lapse series of 10 minutes, 6-7 z-stacks, 1AU pinhole aperture. Image acquisition was performed with ZEN 2.1 software (Carl Zeiss).

A conventional detector was used to acquire the GFP or RFP signal (laser line Argon 25mW and DPSS 10mW, respectively) while the signal from the drug DT-061 was acquired through the laser line Diode 30mW using a highly sensitive multiarray 32PMT GaAsP detector.

The same was performed for iHAP1 for a final concentration of 2µM as a negative control. Stills and videos were further processed using the software Fiji [63].

### Lipidomics

A TNF-sensitive subclone (MCF7-S1) of the MCF7 human ductal breast carcinoma cell line[64](hereafter referred to as MCF7) was cultured in RPMI medium (Gibco) supplemented with 6% FCS and in a humidified incubator set to 37°C and 5% CO_2_. Approximately 1 million MCF7 cells cultured in a 6-well dish were treated with 15 µM DT-061, 1.5 µM iHAP1, 3 µg/mL Brefeldin A, or vehicle (DMSO) for 30 minutes. We then added 10 µM sphingosine (Avanti Polar Lipids, Alabama, USA) or vehicle (DMSO) to the medium and cultured them for additional 4 hours. The cells were then washed five times with ice-cold 155 mM ammonium bicarbonate and harvested into 1 mL 155 mM ammonium bicarbonate by scraping. 200 µL of the cell suspensions were then subjected to lipid extraction and quantitative mass spectrometry-based lipidomics as described previously [65]. The criteria used to identify ceramide, sphingomyelin, hexosylceramide, GM3, and PC are listed in table S4. The internal standards used for quantification of lipids are listed in table S5.

### Statistical analysis

Statistical analysis and graphs were generated in GraphPad Prism 8.0. The data points were tested for normality using Shapiro–Wilk test. Accordingly, statistical significance were determined by unpaired Student’s t-test or Mann-Whitney U test. F test was used to compare variances and for conditions that did not have equal variances, parametric tests with Welch’s correction was used. Details of the statistical significance and n values for each conditions can be found in the figures and figure legends.

### Data availability

The mass spectrometry data have been deposited to ProteomeXchange PXD026666

## Supplementary figure legends

**Figure S1.**
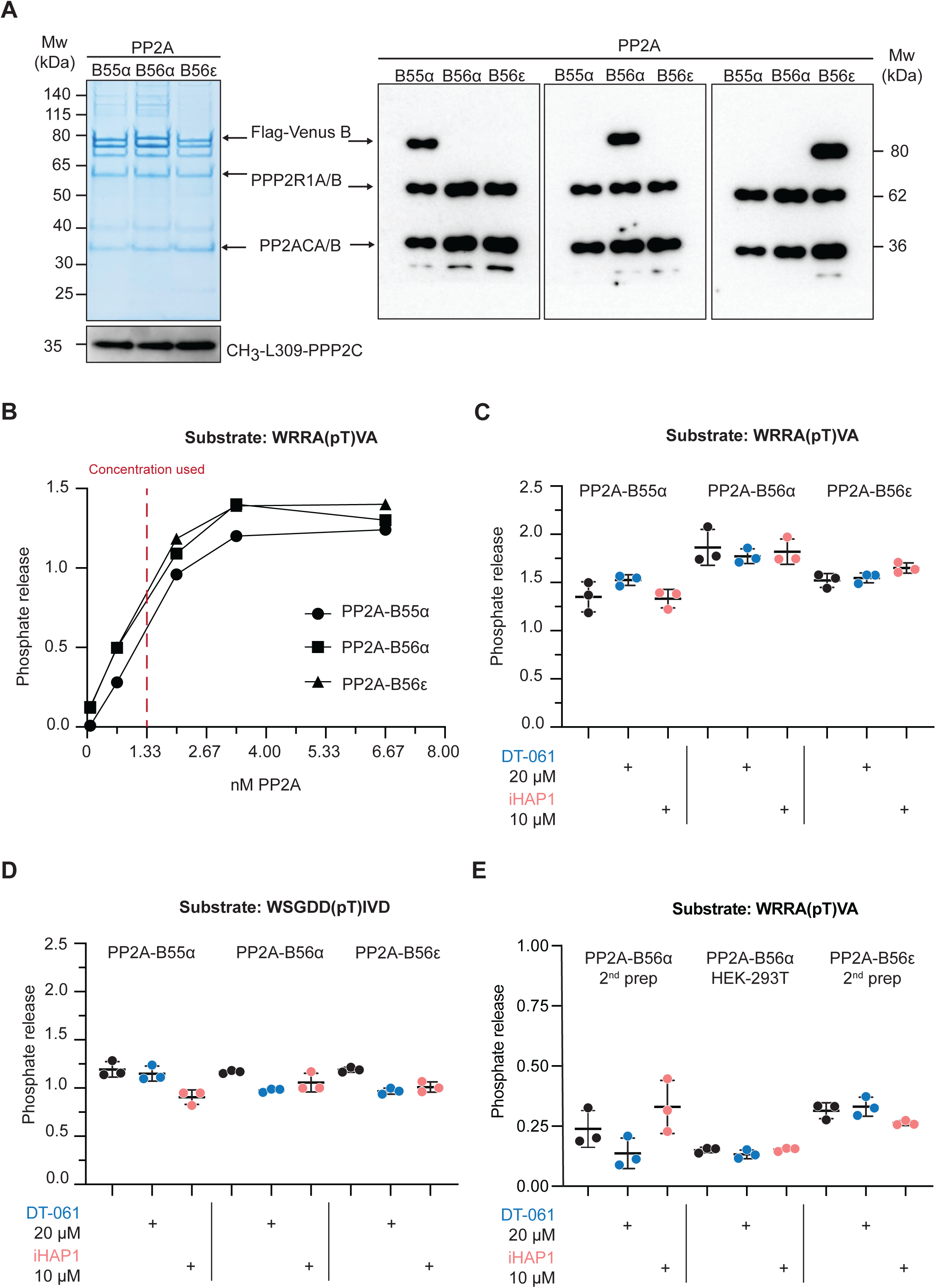
iHAP1 and DT-061 do not activate PP2A complexes *in vitro*. A) Analysis of the purity of PP2A holoenzyme preparations by colloidal staining and Western blotting. Methylation of the C-tail of PPP2C was confirmed using an antibody against CH_3_- L309. Samples were analysed by mass spectrometry which confirmed that no other protein phosphatases were present in the preparations. B) Titration experiment of different PP2A holoenzyme preparations to determine appropriate concentration to use in enzymatic assays. C-D) *In vitro* dephosphorylation assays using model phosphopeptides and purified PP2A holoenzymes. The phosphopeptides were incubated with specific PP2A holoenzymes together with iHAP1 or DT-061 as indicated. The reaction was stopped after 15 minutes, and the amount of phosphate released was measured. Mean and SD from three technical replicates of one out of three independent experiments is shown. E) *In vitro* dephosphorylation assays using a different preparation of PP2A holoenzymes as well as a PP2A-B56α complex reconstituted by expressing all 3 subunits in HEK-293T cells.

**Figure S2.**
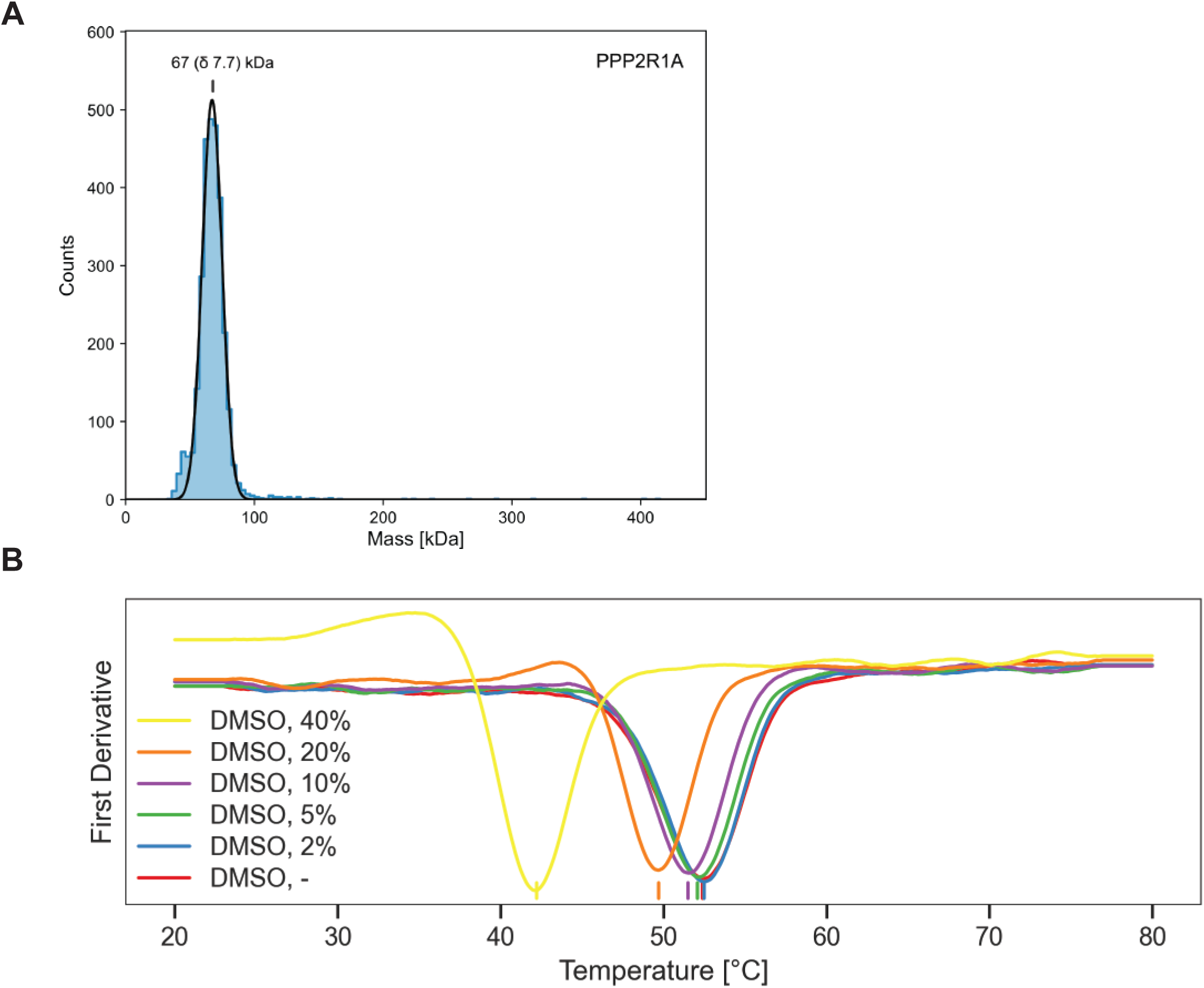
Characterization of PPP2R1A preparation and stability in DMSO. A) MP mass distribution of PPP2R1A in NMR buffer (PBS). The mass photometry measurement of PPP2R1A shows a major peak with molecular weight of 67 kDa in agreement with the molecular weight of the monomeric PPP2R1A. B) First derivative of the integrated F350/F330 nm fluorescence ratio thermal unfolding curves color coded as increasing percentages of DMSO in ITC buffer. T_m_ values of each sample are indicated by vertical lines.

**Figure S3.**
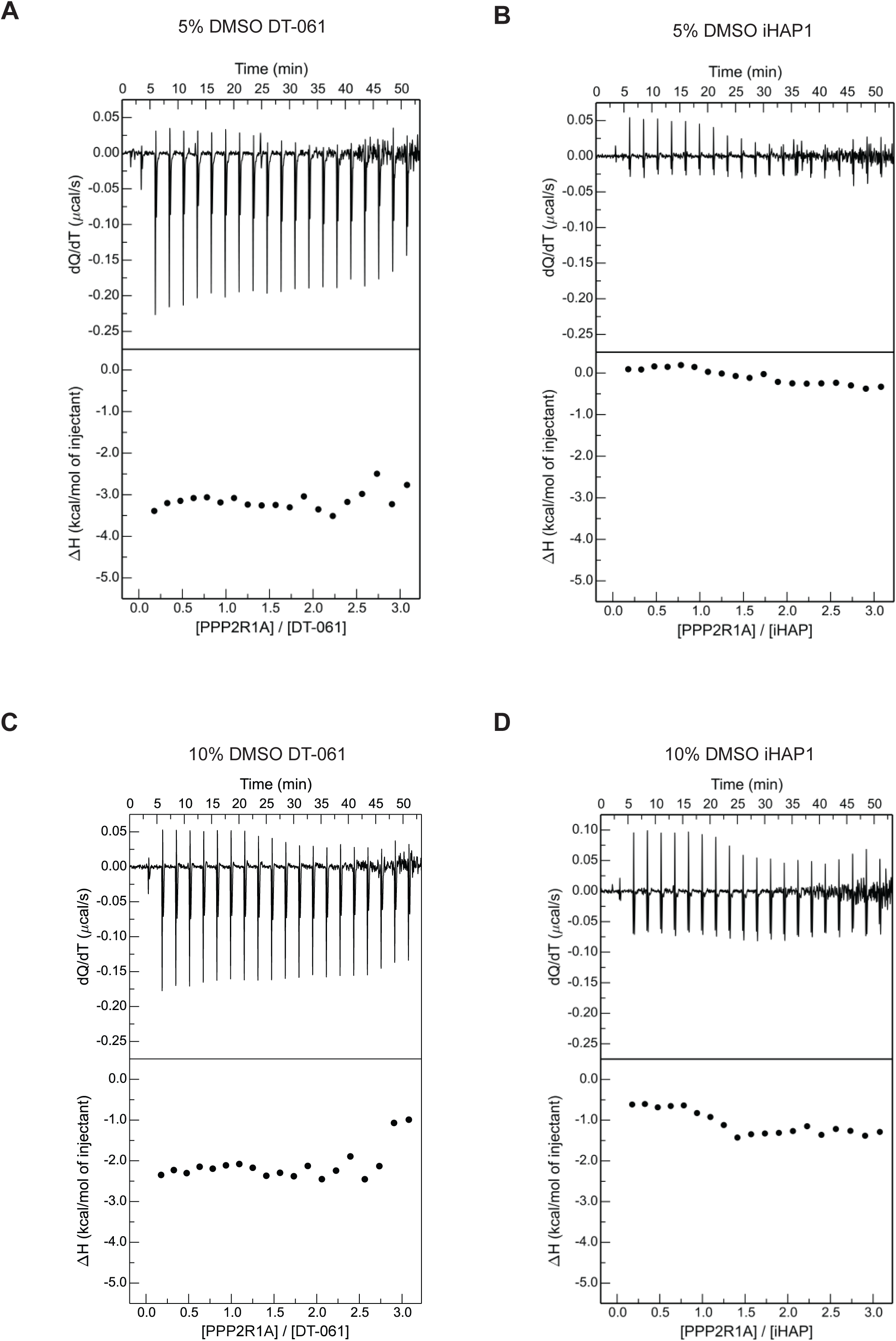
iHAP1 and DT-061 do not bind PPP2R1A. A-D) Thermogram (top panel) for the calorimetric titration of PPP2R1A (300 μM, in syringe) in DT-061 (20 μM, in calorimetric cell) or iHAP (20 μM, in calorimetric cell) at 25 °C and corresponding ligand-normalized integrated heats (bottom panel). Experiments were done in 5% DMSO (A-B) or 10% DMSO (C-D).

**Figure S4.**
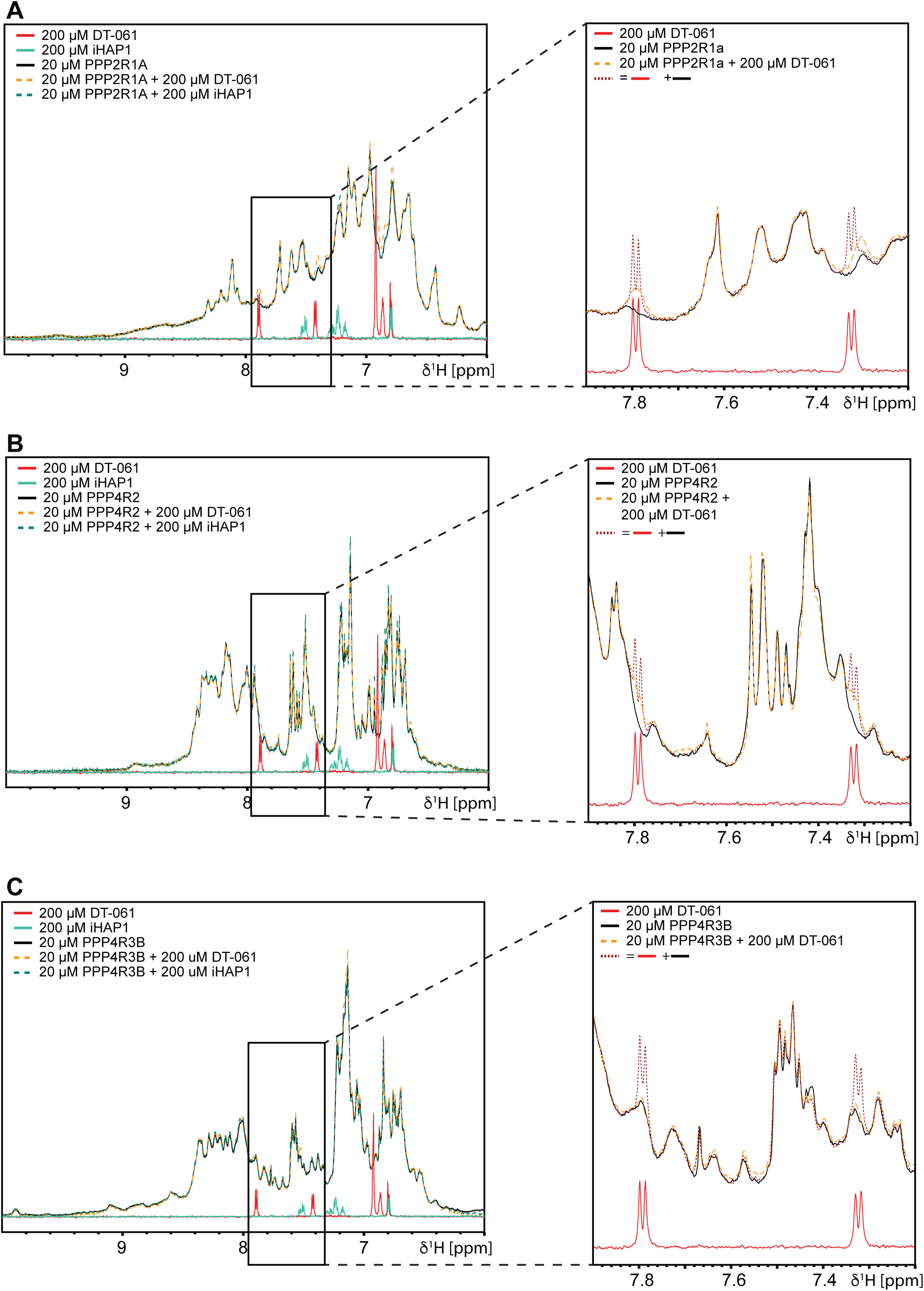
NMR analysis of PPP2R1A interaction with iHAP1 and DT-061. A) Aromatic- and Amide-proton region of the 1D-^1^H-NMR spectra of 20 µM PPP2R1A alone (black) and in presence of 200 µM DT-061 (red dotted) or 200 µM iHAP1 (cyan dotted) respectively. Zoom in on part of the spectrum on the right showing line broadening of DT-061 peaks upon addition of compound. B) Aromatic- and Amide-proton region of the 1D-^1^H-NMR spectra of 20 µM PPP4R2 alone (black) and in presence of 200 µM DT-061 (red dotted) or 200 µM iHAP1 (cyan dotted) respectively. Zoom in on part of the spectrum on the right showing line broadening of DT-061 peaks upon addition of compound. C) Aromatic- and Amide-proton region of the 1D-^1^H-NMR spectra of 20 µM PPP4R3B alone (black) and in presence of 200 µM DT-061 (red dotted) or 200 µM iHAP1 (cyan dotted) respectively. Zoom in on part of the spectrum on the right showing line broadening of DT-061 peaks upon addition of compound.

**Figure S5.**
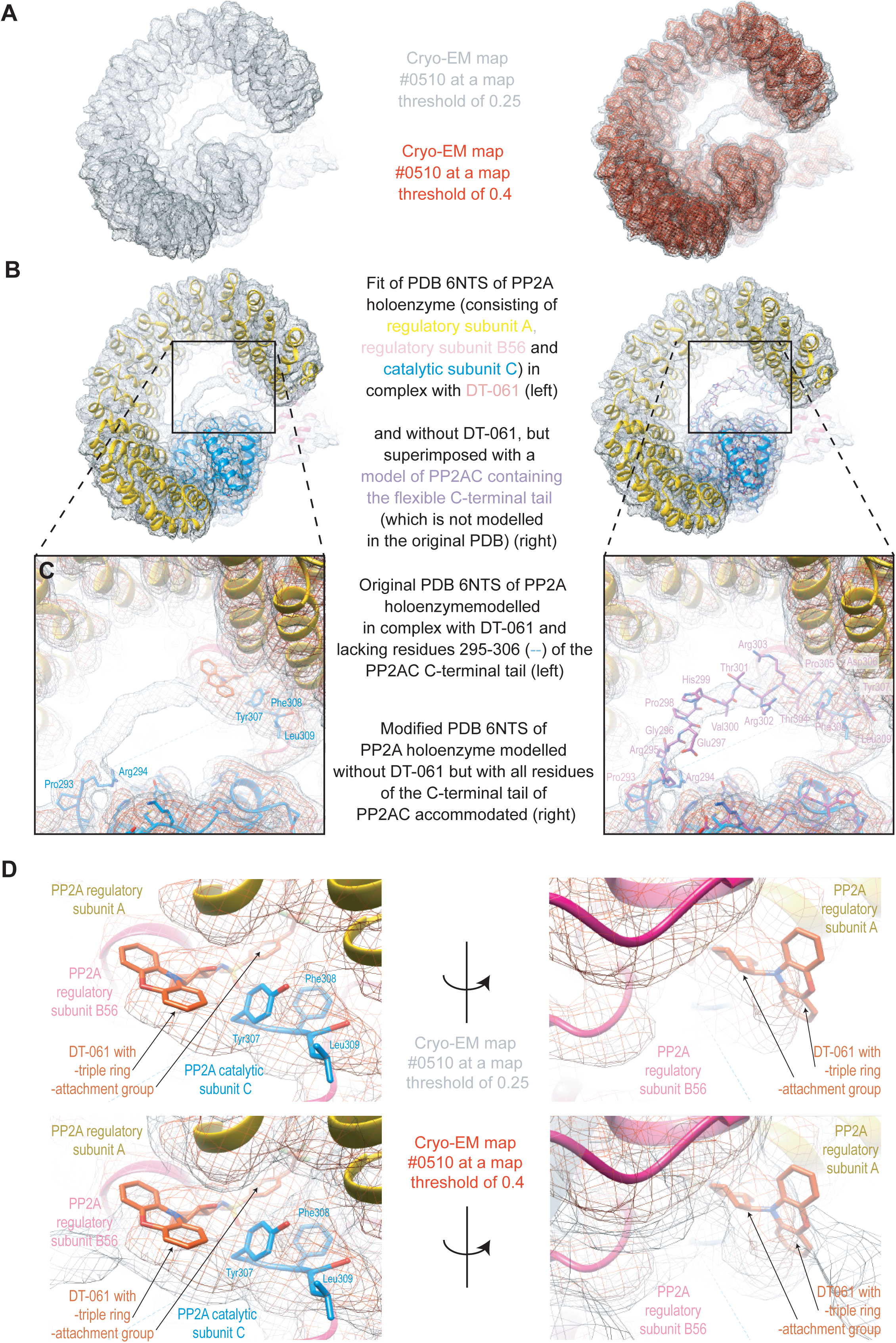
Analysis of the Cryo-EM structure of DT-061 bound to PP2A-B56alpha. Views of EMD-0510 map at two different density threshold levels. Lower threshold levels (0.25, grey) reveal a continuous stretch of extra density which spans the center of the horseshoe-shaped holoenzyme complex. This density is not visible at higher thresholds (0.4, red) indicating that it corresponds to a part of the holoenzyme that is structurally more flexible than the rest. B-C) Fit of PDB 6NTS into the EMD-0510 map, comprising the regulatory A subunit (yellow), the catalytic C subunit (blue) and regulatory B56 subunit (pink). The fit shows that the low threshold extra density connects the main body of the C subunit (modelled residues 1-294, blue) with the three most C-terminal residues of the C subunit (residues 307-309, blue) that are in contact with the regulatory A subunit and in the immediate vicinity of where the DT-061 ligand has been placed (red, left panel). Residues 295-306 are absent from this model of the C subunit (blue dashed line). In the right-hand panel a complete model of the C subunit that contains residues 295-306 (purple, right panel) has been superimposed on the C subunit of PDB 6NTS. The fit of the 295-306 residues is shown for illustrative and comparison purposes. The modeling of that region shows that the presence of the 295-306 residues could also explain the stretch of extra density. The presence of these residues would promote a steric clash with the suggested position of DT-061. D) Detailed view of the suggested DT-061 ligand fit into EMD-0510 map. The two views (left- and right-hand panel) and the two different map thresholds (grey 0.25 and red 0.4, bottom panels) show that the ligand’s attachment group is embedded quite well into the cryo-EM map, but the tricyclic heterocyclic moiety fit appears suboptimal even at lower map thresholds.

**Figure S6.**
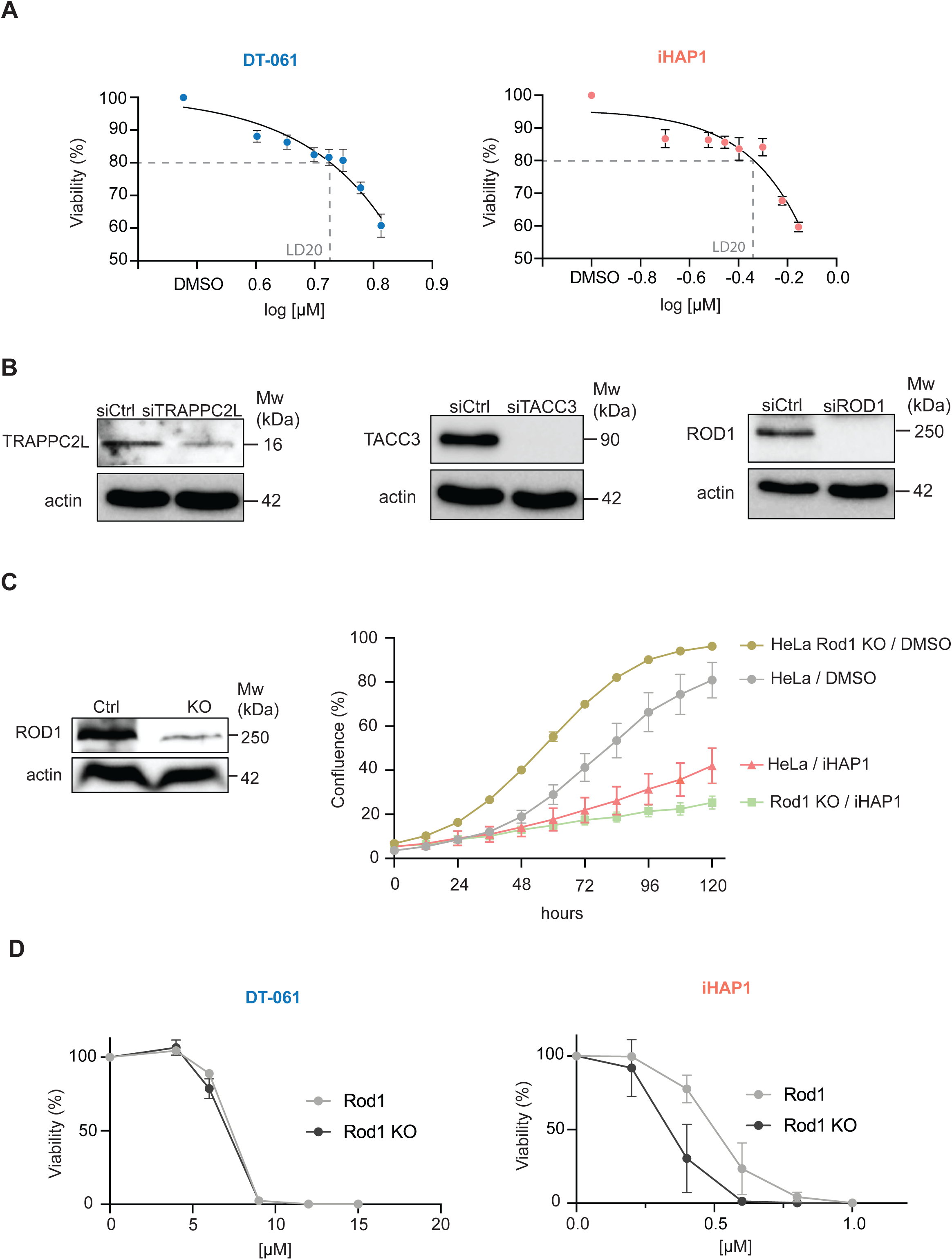
Compound titration and validation of CRISPR screen results. A) Viability of RPE1-hTERT P53-/- FLAG Cas9 cells with increasing concentrations of iHAP1 or DT-061 to determine LD20. B) Protein depletion efficiency after RNAi-treatment. Actin was used as loading control. C) Sensitivity of HeLa Rod1 KO cells to iHAP1. Western blot to confirm Rod1 depletion levels (left) and growth curves for the indicated conditions (right). D) Sensitivity of HeLa Rod1 KO to DT-061 or iHAP1 determined using SRB assay. C-D) Mean and SD of three independent experiments is shown.

**Figure S7.**
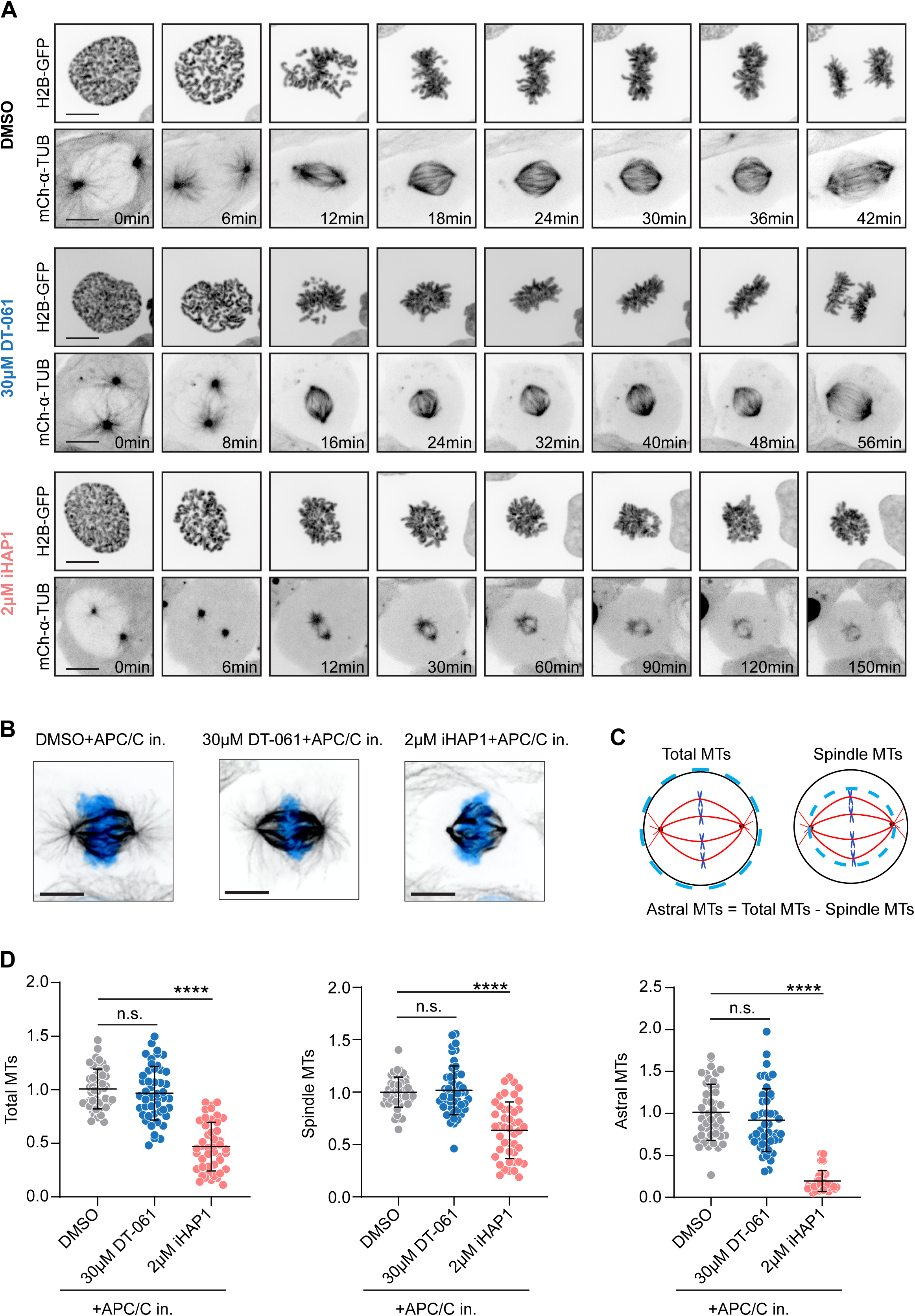
Analysis of mitotic phenotypes in cells treated with DT-061 or iHAP1. A) Representative spinning-disk confocal microscopy time-series of mitosis in U2OS cells stably expressing H2B-GFP/mCherry-α-tubulin following indicated treatments. Scale bar 10µm. B) Representative images of U2OS mitotic spindles in metaphase following indicated treatments, immunostained with α-tubulin antibody. DNA was counterstained with DAPI (cyan). Scale bar 10µm. C) Illustration describing the method used for measuring the amount of total, spindle and astral microtubules. D) Quantification of the number of microtubules in metaphase-arrested mitotic cells following indicated treatments. The mean and SD are plotted from three independent experiments (Total no. of cells = 48 (DMSO), 45 (30 µM DT-061) and, 48 (Ap+pT 2 µM iHAP1)). ns, not significant, ****p<0.0001

**Figure S8.**
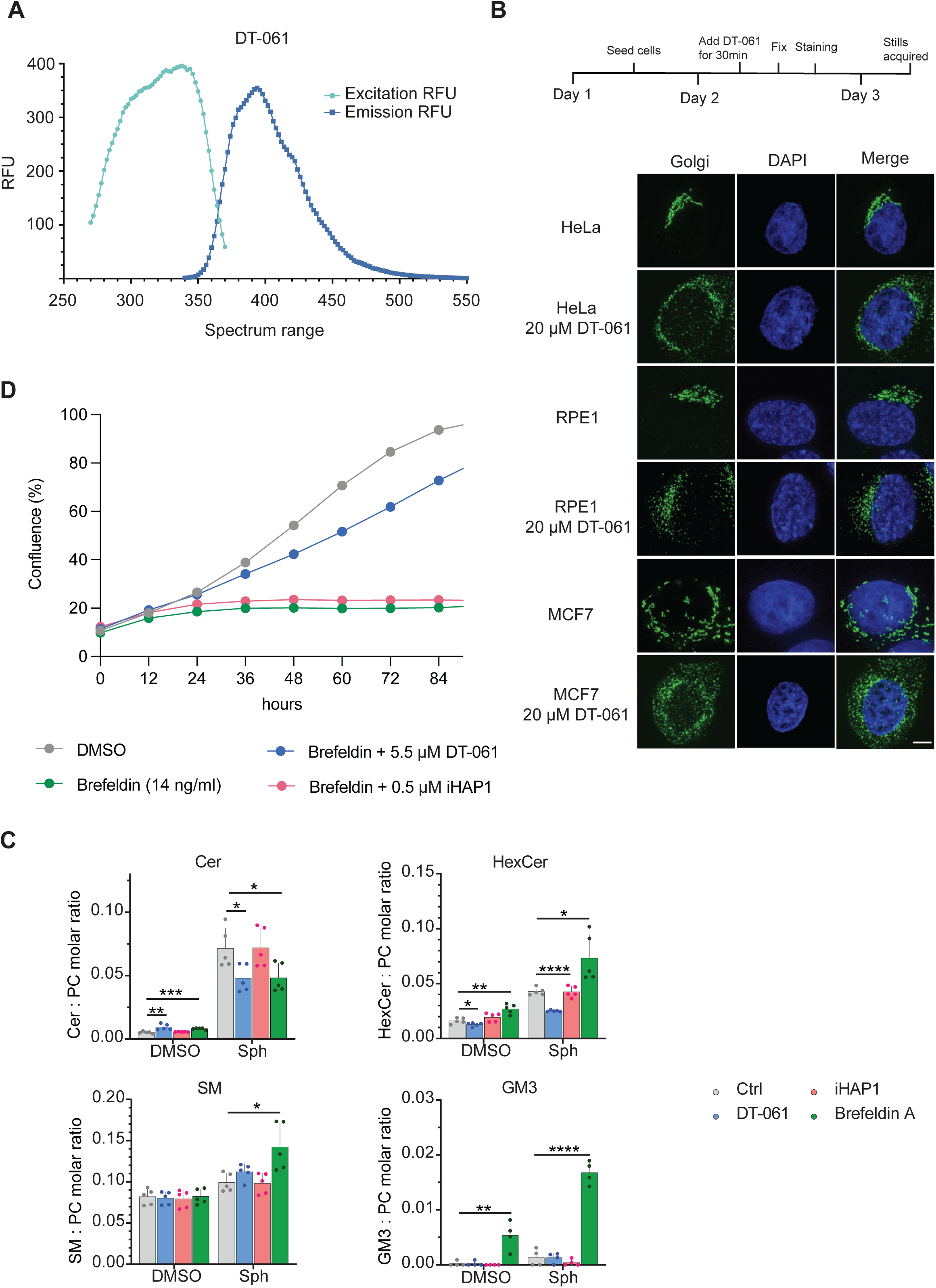
Analysis of DT-061 effects on Golgi. A) Excitation/emission spectrum for DT-061. B) Immunofluorescence analysis of Golgi using an antibody against GM130. The indicated cell lines were treated with DT-061 for 30 minutes before fixation. Representative images shown. Scale bar 5 µm. C) Molar quantities of sphingolipid classes detected in MCF7 cells after feeding with the precursor sphingosine or treatment with vehicle (DMSO). MCF7 cells were co-treated with DT-061, iHAP1, Brefeldin A, or vehicle. The determined molar quantities of sphingolipid classes were normalized to that of PC. The presented values represent averages of five independent experiments. *p<0.05, **p<0.01, ***p<0.001, ****p<0.0001. The abbreviations used are Cer: ceramide, HexCer: hexosylceramide, PC: phosphatidylcholine, and SM: sphingomyelin. D) Growth curves of HeLa cells treated with the indicated combinations of drugs. The results are representative of 4 independent experiments.

## Supplementary tables

Table S1: Mass spectrometry analysis of myc-PP2AC purifications

Table S2: Mass spectrometry analysis of MC-bead assay

Table S3: normZ score for CRISPR screen

Table S4: Lipid identification criteria

Table S5: Internal lipid standards

## Supplementary movie legends

**Movie-1**: Microtubule dynamics visualized in U2OS EB1-GFP cells following indicated treatments.

**Movie-2**: Mitosis in control U2OS cells stably expressing H2B-GFP/mCherry-α-tubulin treated with DMSO. DNA labelled by H2B-GFP on the left panel and microtubules labelled by mCherry-α-tubulin on the right panel.

**Movie-3**: Mitosis in U2OS cells stably expressing H2B-GFP/mCherry-α-tubulin treated with 30µM DT-061 for 20min before filming. DNA labelled by H2B-GFP on the left panel and microtubules labelled by mCherry-α-tubulin on the right panel.

**Movie-4**: Mitosis in U2OS cells stably expressing H2B-GFP/mCherry-α-tubulin treated with 2µM iHAP1 for 20min before filming. DNA labelled by H2B-GFP on the left panel and microtubules labelled by mCherry-α-tubulin on the right panel.

**Movie-5**: Live cell imaging of cell expressing the Golgi marker treated with DT-061 (10 minutes time-lapse).

**Movie-6**: Live cell imaging of cell expressing the ER marker treated with DT-061 (10 minutes time-lapse).

